# Securin Regulates the Spatiotemporal Dynamics of Separase

**DOI:** 10.1101/2023.12.12.571338

**Authors:** Christopher G. Sorensen Turpin, Dillon Sloan, Marian LaForest, Lindsey Uehlein Klebanow, Diana Mitchell, Aaron F. Severson, Joshua N. Bembenek

## Abstract

Separase is a key regulator of the metaphase to anaphase transition with multiple functions. Separase cleaves cohesin to allow chromosome segregation and localizes to vesicles to promote exocytosis in mid-anaphase. The anaphase promoting complex/cyclosome (APC/C) activates separase by ubiquitinating its inhibitory chaperone, securin, triggering its degradation. How this pathway controls the exocytic function of separase has not been investigated. During meiosis I, securin is degraded over several minutes, while separase rapidly relocalizes from kinetochore structures at the spindle and cortex to sites of action on chromosomes and vesicles at anaphase onset. The loss of cohesin coincides with the relocalization of separase to the chromosome midbivalent at anaphase onset. APC/C depletion prevents separase relocalization, while securin depletion causes precocious separase relocalization. Expression of non-degradable securin inhibits chromosome segregation, exocytosis, and separase localization to vesicles but not to the anaphase spindle. We conclude that APC/C mediated securin degradation controls separase localization. This spatiotemporal regulation will impact the effective local concentration of separase for more precise targeting of substrates in anaphase.

## Introduction

The cell cycle is comprised of distinct phases punctuated by checkpoints that function as gatekeepers to control progress through critical events. During mitosis, the metaphase to anaphase transition is regulated by the spindle assembly checkpoint (SAC) to ensure proper chromosome segregation (McAinsh & Kops, 2023; Musacchio, 2015). In prometaphase, kinetochores on chromosomes capture spindle microtubules from opposite poles to achieve bioriented spindle attachment in metaphase (Foley & Kapoor, 2013). The SAC surveilles for proper kinetochore attachment to the spindle and delays anaphase onset until all chromosomes are properly aligned on the metaphase plate (Lara-Gonzalez et al., 2021). Once chromosomes are properly aligned, SAC signaling is silenced and anaphase commences. This checkpoint ensures that each daughter cell inherits a complete set of chromosomes.

Entry into anaphase requires an E3 ubiquitin ligase called the anaphase promoting complex/cyclosome (APC/C; Alfieri et al., 2017). APC/C targets multiple substrates for polyubiquitination, which designates them for degradation by the 26S proteosome (Watson et al., 2019). In metaphase, APC/C is kept inactive by the SAC. Once the checkpoint is satisfied, the APC/C ubiquitinates securin, an inhibitory chaperone of separase. Separase is the protease that cleaves a subunit of cohesin, the molecular glue that holds chromosomes together after their duplication in S-phase (Uhlmann et al., 2000). The cleavage of cohesin by separase allows the poleward movement of chromosomes, the defining event of anaphase onset, and is therefore a critical regulatory mechanism for successful cell division (Uhlmann, 2003).

This regulatory pathway also controls chromosome segregation during meiosis to facilitate the formation of healthy gametes (Gorbsky, 2015). Oocyte meiosis consists of two rounds of chromosome segregation and polar body extrusion to create a single large gamete (Mogessie et al., 2018). In many species, oocytes arrest in metaphase II prior to fertilization, in part due to inhibition of APC/C (Tunquist & Maller, 2003). Fertilization triggers a process called egg activation that transforms the arrested oocyte into a rapidly dividing embryo. Egg activation events include the metaphase to anaphase transition, which requires separase mediated removal of cohesin. In addition, egg activation triggers the exocytosis of special vesicles called cortical granules (Horner & Wolfner, 2008). Exocytosis of cortical granule cargo modifies the extracellular matrix of the oocyte to block polyspermy, and ensures a mechanically stable, chemically uniform environment for early development (Liu, 2011; Liu et al., 2003; Wessel et al., 2001). Although highly specialized, many fundamental cell cycle discoveries have been made by studying oocyte meiosis.

Many questions remain about the regulation of egg activation, in part owing to the difficulty of conducting *in vivo* analysis of female meiosis in many models (Robker et al., 2018). *C. elegans* allows for the observation of the entirety of female meiosis *in vivo* (Lui & Colaiácovo, 2013). Egg activation occurs during meiosis I in *C. elegans* oocytes just after fertilization and depends on sperm signals (McCarter et al., 1999; McNally & McNally, 2005; Yang et al., 2005). Cortical granule exocytosis occurs during anaphase I, releasing cargo to build a complex extracellular matrix called the eggshell (Bembenek et al., 2007; Olson et al., 2012). Depletion of several cell cycle regulators, including the APC/C, securin and separase, disrupts eggshell formation (Bembenek et al., 2007; Shakes et al., 2003; Sönnichsen et al., 2005). Importantly, separase localizes to cortical granules in anaphase I and is required for their exocytosis (Bembenek et al., 2007). The protease activity of separase is required for exocytosis of RAB-11 positive vesicles during both meiotic and mitotic anaphase (Bai & Bembenek, 2017; Bembenek et al., 2010; Mitchell et al., 2014). These findings implicate metaphase to anaphase transition regulators in vesicle trafficking. How the SAC pathway regulates separase function in exocytosis has not been investigated.

Regulators of the metaphase to anaphase transition have been implicated in the control of membrane trafficking (Bembenek et al., 2007; Sönnichsen et al., 2005). In *Drosophila*, a component of the RZZ kinetochore complex involved in SAC signaling is involved in Golgi-related trafficking (Wainman et al., 2012). In human cells, securin is known as pituitary tumor transforming gene, is overexpressed in cancers and causes altered secretion (Donangelo et al., 2006; Heaney et al., 1999; Yu et al., 2000). Securin and separase also affect membrane traffic in mammalian and plant cells (Bacac et al., 2011; Moschou et al., 2013). The regulation of membrane trafficking during cell division is poorly understood, and how the cell cycle machinery controls the chromosome and vesicle functions of separase is unknown.

The activation of separase through securin degradation has been extensively characterized using biochemical methods. However, the spatiotemporal dynamics of separase subcellular localization are not well characterized. In yeast, securin is required for proper nuclear and spindle localization of separase (Agarwal & Cohen-Fix, 2002; Jensen et al., 2001). Determining separase localization in humans cells is difficult for technical reasons, and separase has only been observed on chromosomes when staining over-expressed, tagged separase on chromosome spreads (Chestukhin et al., 2003; Sun et al., 2009). It is possible that fluorescently tagging separase could affect its proper function and/or localization. Various separase biosensors have been used to indirectly observe separase activity at specific subcellular compartments (Agircan & Schiebel, 2014; MacKenzie et al., 2023; Monen et al., 2015; Nam & van Deursen, 2014; Rosen et al., 2019; Weber et al., 2020). The biosensor chosen will affect whether different intracellular sites can be monitored, and a biosensor for vesicles has not been studied. In *C. elegans*, separase localization has been verified with both antibody staining and GFP tagging, allowing for direct and accurate spatiotemporal analysis (Bembenek *et al.,* 2007). Therefore, *C. elegans* female meiosis is an ideal context to investigate how separase localization is regulated.

In *C. elegans* meiosis, separase localizes at kinetochore cup structures in prometaphase I and then appears at the midbivalent, where the meiotic cohesin that is destroyed in anaphase I is located (Bembenek et al., 2007; Muscat et al., 2015; Severson & Meyer, 2014). The precise timing and regulation of separase dynamics have not been investigated. In metaphase, separase is present in cortical “linear elements,” where numerous kinetochore proteins also localize. Separase then appears on vesicles in anaphase, where it likely cleaves a target to promote exocytosis (Bai & Bembenek, 2017; Bembenek et al., 2007). Whether the canonical spindle checkpoint pathway regulates the localization of separase is unknown. Previously, it was reported that securin is degraded over a several minute period around the time of anaphase onset (Wang et al., 2013). Whether securin degradation is required for separase activity on vesicles in an important open question. In addition, chromosome segregation begins prior to exocytosis and securin degradation might control the timing of different separase dependent events. Understanding how separase reaches sites of action and how the cell cycle machinery controls this is important to fully understand how separase performs its cellular functions.

To address these questions, we used live imaging with high spatiotemporal resolution to compare separase dynamics during meiosis I to the dynamics of securin degradation and cohesin loss. While securin is degraded over several minutes, separase undergoes rapid localization dynamics seconds before chromosomes move apart. Cohesin is removed from chromosomes within seconds of separase relocalization, initiating the poleward movement of chromosomes at anaphase onset. We investigated how depletion of securin and the APC/C affect chromosome segregation, vesicle trafficking and separase localization. Securin and APC/C depletion causes pleiotropic cell cycle defects. Therefore, we expressed a non-degradable mutant of securin to determine how securin affects separase activity and localization in cells with normal APC/C activity. Our findings suggest a novel role for the APC/C and securin in controlling the localization of separase, in addition to their known role in regulating separase protease activity for chromosome segregation and exocytosis during anaphase I.

## Results

### Securin and Separase Dynamics during Meiosis I

While securin degradation begins prior to anaphase onset (Wang et al., 2013), it is not clear whether securin is degraded before or after separase localizes to sites of action in anaphase. To address this question, we generated worms with endogenously tagged separase (SEP-1::GFP) or securin (IFY-1::GFP) with the chromosome marker H2B::mCherry to assess their localization during meiosis I. As shown previously, SEP-1::GFP localization is dynamic during meiosis I (Fig. 1, A-D; Bembenek *et al.,* 2007). During prophase I, SEP-1::GFP is cytoplasmic and excluded from the nucleus (Fig. 1 A). Just before nuclear envelope breakdown (NEBD), SEP-1::GFP accumulates in the nucleus and becomes enriched on kinetochores (Fig. 1 B).

**Figure 1.**
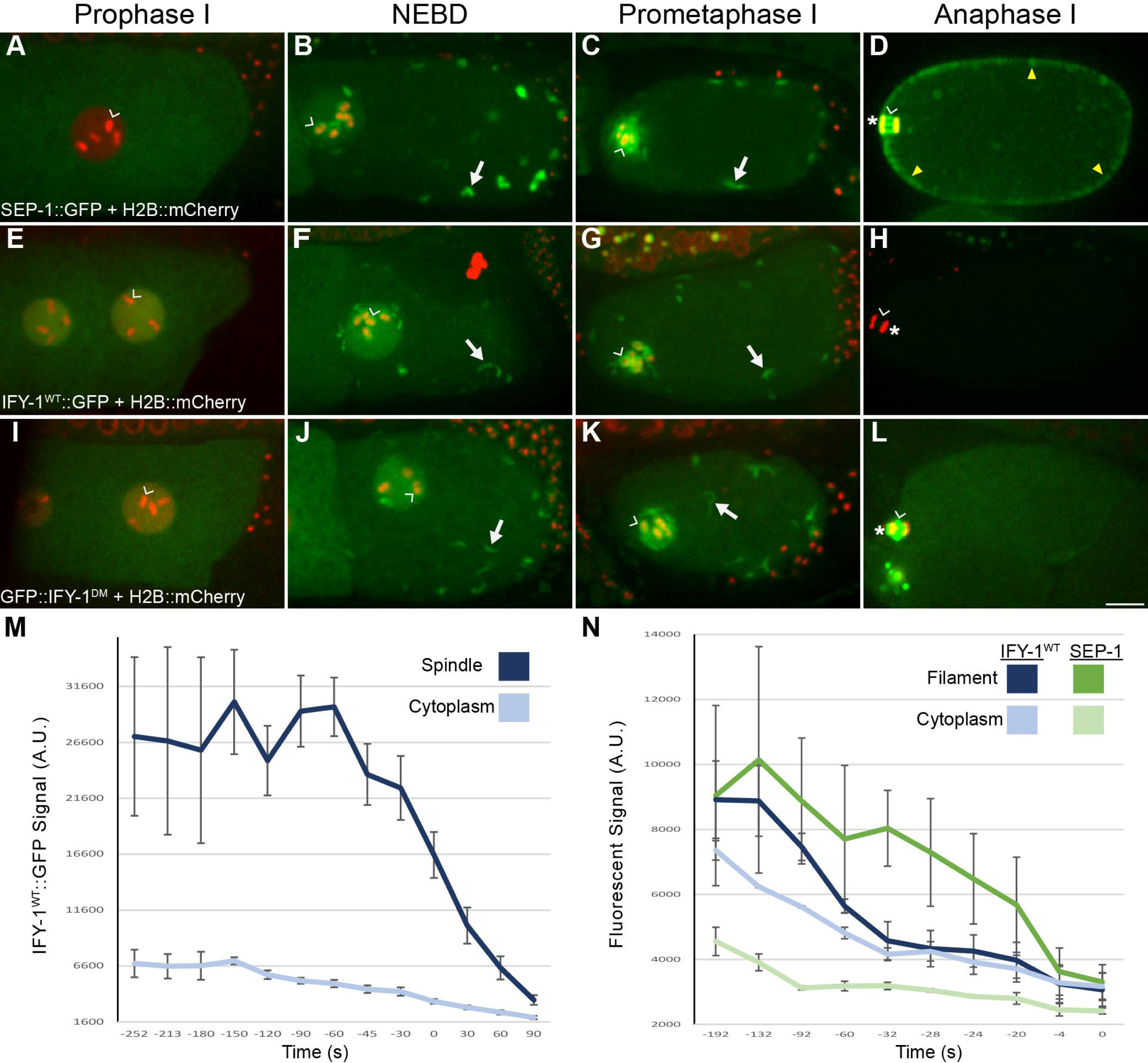
Separase and Securin dynamics in meiosis I. (A-D) Representative images SEP-1::GFP (green) with H2B::mCherry (red) during meiosis I. (A) During prophase I, SEP-1::GFP is cytoplasmic and excluded from the nucleus. (B) At NEBD, SEP-1::GFP accumulates in the nucleus, on chromosomes (caret) and to cytoplasmic kinetochore linear elements throughout the cortex (white arrows), where it remains throughout prometaphase (C). (D) By mid-anaphase, separase localizes between separating chromosomes (caret) and to vesicles (arrowheads). (E-L) Embryos expressing H2B::mCherry (red) with (E-H) IFY-1^WT^::GFP (green) or (I-L) IFY-1^DM^::GFP (green) in meiosis I. (E) IFY-1^WT^::GFP and (I) IFY-1^DM^::GFP are present in both the cytoplasm and the nucleus in prophase oocytes. IFY-1^WT^::GFP (F) and IFY-1^DM^::GFP (J) display identical localization patterns as separase at NEBD and through prometaphase I (G, K). In anaphase I, IFY-1^WT^::GFP (H) is mostly degraded and is not observed on vesicles. In contrast, IFY-1^DM^::GFP (L) is not degraded in anaphase and accumulates on chromosomes (caret) and the anaphase I spindle (white arrowhead) but does not enrich on cortical granules. (M) Quantification of IFY-1^WT^::GFP spindle-associated and cytoplasmic signal showing rapid degradation (t = 0 is chromosome separation at anaphase onset). (N) Quantification of endogenously tagged SEP-1::mCherry and IFY-1^WT^::GFP localized to linear elements in the cortex, (t = 0 is separase localization to vesicles). Securin levels equilibrate with cytoplasmic signal before separase leaves linear elements and appears on vesicles. Scale bar: 10µm.

Simultaneously, SEP-1::GFP appears on cytoplasmic “linear element” structures throughout the cortex and around the nucleus (Fig. 1 B-C). We previously found that separase colocalizes with outer kinetochore proteins on chromosomes and linear elements in prometaphase I (Bembenek et al., 2007). In anaphase, SEP-1::GFP remains on the spindle and chromosomes but no longer appears on linear elements in the cortex and is instead found on vesicles until they undergo exocytosis (Fig. 1 D). How separase localization to these various subcellular sites is regulated is not known.

To learn more about separase regulation, we investigated securin (IFY-1) localization during meiosis I. During prophase I, IFY-1 is detected in the cytoplasm and is enriched in the nucleus at the time when separase is excluded from the nucleus (Fig. 1 E). This nuclear pool of IFY-1 is therefore likely not bound to separase. An excess pool of securin is also found in mammalian oocytes and serves as a competitive substrate of APC/C to time cell cycle events (Thomas et al., 2021). At NEBD through prometaphase I, IFY-1 localizes similarly to SEP-1 at kinetochore cups on the chromosomes, and linear elements in the cortex (Fig. 1 F-G). This indicates that separase is likely securin-bound and thus inactive when it localizes to kinetochore structures, as expected from previous studies. IFY-1 signal begins to decline in the cytoplasm by late prometaphase I, approximately three minutes before anaphase onset, as previously described in worms (Fig. 1 M, Wang et al., 2013). At anaphase onset, significantly reduced IFY-1 signal can be detected at the central spindle, which rapidly lowers to cytoplasmic levels (Fig. 1 H, M). Importantly we did not detect IFY-1 on vesicles at any timepoint (Fig. 1 E-H). Endogenously tagged and multiple, independent transgenic IFY-1 GFP lines show similar degradation curves regardless of expression level (Fig. S1 E). In the cortex, IFY-1::GFP and SEP-1::mScarlet co-localize on linear elements until anaphase I when SEP-1::mScarlet localizes to vesicles and IFY-1::GFP is degraded (Fig. S 1). Therefore, SEP-1 colocalizes with IFY-1 at kinetochore structures until anaphase when IFY-1 is degraded and SEP-1 localizes to sites of action on chromosomes and vesicles. These observations suggest that IFY-1 degradation may be required for SEP-1 to localize to sites of action during anaphase I.

### Separase moves from kinetochore structures to sites of action at anaphase onset

Anaphase onset is defined as the moment when chromosomes move poleward. To better define the dynamics of anaphase onset, we imaged separase, securin and cohesin at a high temporal resolution at the metaphase to anaphase transition. This analysis revealed a previously undocumented, rapid change in separase localization within seconds of anaphase onset (Fig. 2 A-C). During prometaphase I, separase localizes to kinetochore cups on the chromosomes (Fig. 2 A) but is excluded from the midbivalent region where the meiotic cohesin kleisin subunit, GFP::COH-3, localizes (Fig. 2 A-C). After spindle rotation, separase enriches at spindle poles approximately 30 seconds before anaphase I onset (Fig. 2 A-C). Within 10 seconds before anaphase onset, separase begins to invade the midbivalent region of the homologous chromosomes, colocalizing with GFP::COH-3 for a brief period (Fig. 2 A-C). When chromosomes begin to move poleward at anaphase onset, separase is highly accumulated at the midbivalent region and GFP::COH-3 is lost (Fig. 2 A-C, S2 A). Therefore, separase is spatially restricted from the region where cohesin resides until seconds prior to anaphase onset, which could be an important mechanism to precisely time cohesin cleavage.

**Figure 2.**
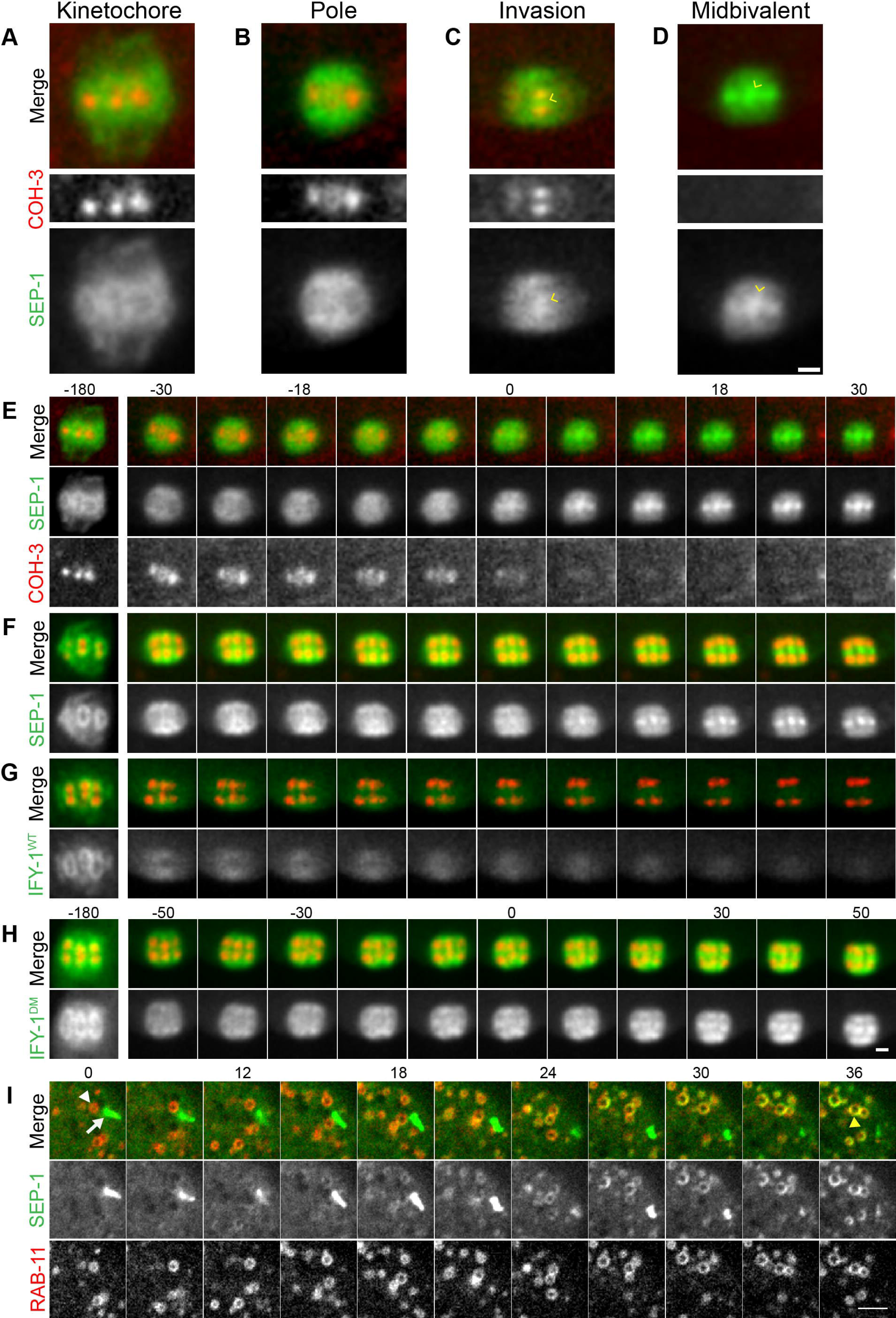
Spatiotemporal dynamics of Separase, Securin and Cohesin at Anaphase Onset. (A-B) Representative localization pattern of SEP-1::mScarlet (green) co-expressed with cohesin subunit COH-3::GFP (red) at the spindle in anaphase I. (A) In prometaphase I, spindle SEP-1::mScarlet localizes to kinetochore cups around the bivalents and is spatially isolated from COH-3::GFP at the midbivalent. (B) Just before anaphase onset, SEP-1::mScarlet accumulates on spindle poles, still not enriched with COH-3::GFP. (C) Seconds before chromosome segregation, SEP-1::mScarlet invades the midbivalent, colocalizing briefly with COH-3::GFP (caret). (D) COH-3::GFP is quickly lost from chromosomes right after SEP-1::mScarlet invades the midbivalent region and chromosomes immediately move poleward at anaphase onset (caret). (E) Montage showing rapid SEP-1::mScarlet localization changes on the spindle and abrupt loss of COH-3::GFP at anaphase onset (time is shown in seconds relative to t=0, which indicates the first timepoint with significant midbivalent accumulation of SEP-1::mScarlet). (F-H) Montages showing the dynamics of SEP-1::GFP, IFY-1^WT^::GFP and GFP::IFY-1^DM^ (green) co-expressed with chromosome marker H2B::mCherry (red) during the metaphase-to-anaphase I transition (t=0 marks when chromosome move apart in F, G, or when midbivalent signal appears in H). (F) SEP-1::GFP appears prominently at the midbivalent when chromosomes move apart at anaphase onset. (G) IFY-1^WT^::GFP signal is significantly reduced by anaphase onset. (H) GFP::IFY-1^DM^ remains high on the spindle throughout anaphase I and localizes similar to separase. (I) In the cortex, SEP-1::GFP relocalizes from linear elements (white arrow) to cortical granules labeled with mScarlet::RAB-11.1 (arrowhead) by 30 seconds after anaphase onset. Scale bars: 2µm.

It is well established that SEP-1 is catalytically activated when IFY-1 is degraded. The relative timing of IFY-1 degradation and SEP-1 movement and localization to sites of action has not been characterized. To address this, we imaged GFP::IFY-1 with H2B::mCherry at high temporal resolution. From NEBD through prometaphase I, the levels of chromosome and spindle associated IFY-1::GFP remains relatively constant (Fig. 1M, Fig. 2D). During early metaphase I, the APC/C is required for the meiotic spindle to become shorter and more compact, rotate, and move into close proximity with the cortex (Yang et al., 2003). Spindle associated IFY-1::GFP gets redistributed and concentrated in the same pattern as separase, but the signal drops precipitously at spindle rotation, losing approximately a third of its average intensity by anaphase I onset (Fig. 2 D, Fig. 1 M, N, Fig. S 1). During the rapid relocalization of separase from kinetochores to the midbivalent, securin levels are rapidly decreasing (Fig. 2 D; Fig. 1 N). In the cortex, SEP-1 and IFY-1 are recruited to linear elements just before NEBD and remain there throughout prometaphase I (Fig. 1B, C, F, G). We imaged endogenously tagged SEP-1::GFP with an endogenously tagged mScarlet::RAB-11 to label cortical granules (Fig. 2 F). After anaphase onset, SEP-1 signal is lost from linear elements and rapidly accumulates on cortical granules within 30 seconds (Fig. 2 F). At the cortical kinetochore filaments, IFY-1::GFP drops to near cytoplasmic levels before SEP-1::mScarlet is lost from the filaments and appears on vesicles (Fig. 1 N). These results indicate that securin levels decrease more rapidly from kinetochore structures than the gradual loss in the cytoplasm. Therefore, securin degradation is rapidly occurring when separase moves from kinetochore structures to the midbivalent and vesicles at anaphase onset.

### The APC/C-pathway controls SEP-1 localization in meiosis I

The rapid degradation of securin, which begins immediately before separase localizes to sites of action, suggests that it may regulate separase localization in addition to its protease activity. If this were true, then depletion of IFY-1 by RNAi might lead to premature loss of securin and aberrant regulation of separase localization. A previous report indicated that SEP-1 does not localize to cortical granules after *ify-1* RNAi in *C. elegans* (Kimura & Kimura, 2012). We repeated this experiment to observe separase localization to chromosomes and vesicles. In control animals, the cortical granule cargo protein mCherry::CPG-2 is observed in vesicles in oocytes and embryos that have not completed exocytosis in anaphase I, while older embryos have mCherry::CPG-2 in the eggshell (Fig. 3 A). In contrast to the previous report, we detected vesicle localization of separase in multiple embryos in the uterus of *ify-1(RNAi)* treated animals (N=16/30 animals, Fig. 3 B). To further validate this result, we characterized separase localization in animals with different RNAi penetrance expressing SEP-1::GFP together with mCherry fused to H2B and CPG-2 to mark chromosomes and vesicles. In mild phenotype cases (14-48 hours of feeding RNAi), vesicle localization of separase is only observed in one or fewer embryos, and most embryos have normal mCherry::CPG-2 in the eggshell, with some polar body extrusion defects and later cell division failures (N=31/101 animals). Animals with an intermediate phenotype (16-39 hours of feeding) contained 2-3 unicellular embryos with SEP-1::GFP localized to mCherry::CPG-2 vesicles, while older embryos had increasing amounts of eggshell mCherry::CPG-2 signal, suggesting reduced and delayed exocytosis of cortical granules (N=34/95, Fig. 3 B). In animals with a severe phenotype (24-48 hours of feeding RNAi), all embryos were unicellular, lacked extracellular mCherry::CPG-2 signal, and had cortical granules trapped in the cytoplasm (N=31/61, Fig. 3 C). In severe *ify-1(RNAi)* embryos, SEP-1::GFP signal in the spindle region and at cortical granules is reduced, but can still be observed. Therefore, securin depletion does not prevent separase localization to vesicles, but inhibits exocytosis.

**Figure 3.**
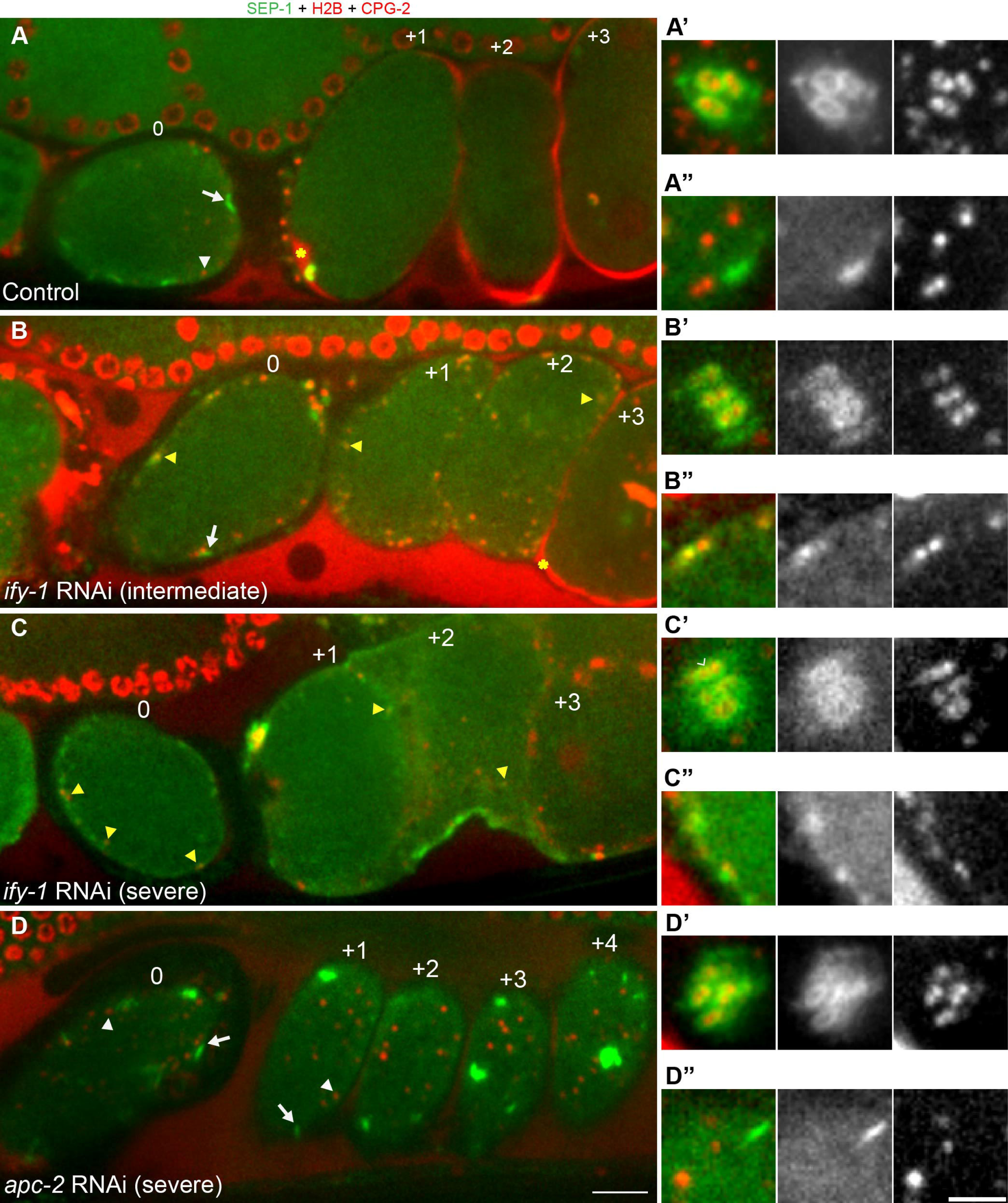
The APC/C and securin control SEP-1 localization in meiosis I. Images of the germline in worms expressing SEP-1:::GFP (green) with chromosome marker H2B::mCherry and cortical granule cargo protein mCherry::CPG-2 (red). Insets show high magnification of the spindle in A’-D’ or vesicles in A”-D”. Numbers correspond to relative position of embryos in the uterus, which also corresponds to their increasing age as each ovulation occurs every 25-30 minutes. (A) In control animals fed OP50, prometaphase I embryos (embryo 0) have mCherry::CPG-2 in vesicles, while older embryos (+1 to +3) have mCherry::CPG-2 incorporated into the eggshell (denoted by yellow asterisk in the +1 embryo). Prometaphase I embryos in the spermatheca have SEP-1::GFP localized to kinetochores at the spindle (A’) and linear elements in the cortex (A”). (B) Intermediate *ify-1* RNAi animals have prometaphase I embryos in the spermatheca (0) with SEP-1::GFP localized abnormally to chromosomes in the midbivalent region (B’) and localization to vesicles (yellow arrowhead in C, cortical region in B”). Multiple older embryos (+1-+2) also have SEP-1::GFP localized to cortical granules (yellow arrowheads). Early embryos lack eggshell signal but eventually some mCherry::CPG-2 signal appears outside older embryos (+3, asterisk). (C) Severe *ify-1* RNAi causes SEP-1::GFP to remain on vesicles in older embryos (+1, +2, yellow arrowheads). mCherry::CPG-2 remains trapped in cortical granules and no eggshell is detected in any of the embryos at any age. SEP-1::GFP displays mislocalized chromosome (C’, caret denotes signal at midbivalent) and vesicle (C”) localization in prometaphase I embryos. (D) After *apc/c* RNAi, separase localization is normal in prometaphase I embryos in the spermatheca (embryo 0), appearing on the kinetochore cups at the spindle (B’) and cortical linear elements (B”). Arrested embryos (+1-+4) in the uterus all show SEP-1:::GFP on kinetochore structures (white arrows). mCherry::CPG-2 remains in cortical granules (white arrowheads) and does not incorporate into the eggshell. Scale bars: 10µm for A-D; 5µm for insets.

We also depleted a negative regulator of securin, the APC/C, and examined separase localization. APC/C was also found to be required for separase to localize to vesicles (Kimura & Kimura, 2012). In one cell arrested *apc-2(RNAi)* embryos, cortical granule exocytosis does not occur and embryos never display mCherry::CPG-2 signal in the extracellular space (Fig. 3 D). As previously shown, SEP-1 does not appear on vesicles (Fig. 3 D), but remains localized to kinetochore cups and linear elements (Fig. 3 D’ and D”). As expected, *apc-2* RNAi blocks degradation of IFY-1::GFP, which remains trapped with SEP-1::mCherry on kinetochore structures in arrested embryos (Fig. S 2). Therefore, APC/C is required for securin degradation and separase localization to vesicles, causing separase to remain associated with kinetochore structures at the spindle and cortex.

One prediction for APC/C-mediated spatiotemporal regulation of separase, via securin destruction, is that depletion of securin should cause separase localization to change prematurely on chromosomes and vesicles. To test this, we examined separase localization in early stages of meiosis by acquiring z-stacks of prometaphase I stage embryos in the spermatheca and by taking time lapse movies from ovulation to anaphase I. In control animals, prometaphase I embryos have SEP-1::GFP at kinetochore cups (Fig. 3 A’) and kinetochore filaments (N=5/5, Fig. 3 A”), with no vesicle localization. Interestingly, in intermediate and severe *ify-1(RNAi)* embryos, separase was mislocalized on chromosomes and could be observed between homologs where cohesin resides in embryos within the spermatheca (N=22/24, Fig 3 B’, C’). In addition, we observed SEP-1::GFP on cortical granules prematurely in *ify-1(RNAi)* embryos within the spermatheca (N=26/35, Fig. 3 B”, C”). *apc-2(RNAi)* embryos in the spermatheca show the same separase localization to kinetochore structures as control (N=5/5, Fig. 3D’, D”). Therefore, securin depletion causes premature relocalization of separase at chromosomes and vesicles, while depletion of the APC/C prevents separase relocalization, consistent with the hypothesis that securin degradation regulates separase spatiotemporal dynamics.

Given that securin depletion causes premature localization of separase to its sites of action, we expected that it may also become proteolytically activated prematurely. We tested this hypothesis in the context of cohesin proteolysis using worms co-expressing COH-3::GFP with chromosome marker H2B::mCherry (Fig. S 3). In WT embryos, COH-3::GFP remains on chromosomes until seconds before chromosome segregation begins (Fig. S3 A). In prometaphase I embryos within the spermatheca, COH-3::GFP signal remains similar in *control* or *apc-2* RNAi (Fig. S3 B, C, E; N=5). In contrast, prometaphase I *ify-1(RNAi)* embryos in the spermatheca had a significant decrease in COH-3::GFP levels on chromosomes (N=4). Therefore, securin depletion causes premature activation of cohesin cleavage on chromosomes and premature relocalization of separase at chromosomes and vesicles.

### Non-degradable Securin expression is dominant negative in *C. elegans*

Inactivation of APC/C and securin have pleiotropic effects on the cell cycle that might impact separase. APC/C has many substrates besides securin, and securin has multiple regulatory functions. To more precisely test whether stabilized securin affects separase localization in anaphase, we expressed a non-degradable IFY-1 mutant from a transgene in a WT background with endogenous IFY-1 and characterized the phenotype. The unstructured IFY-1 N-terminus has a conserved destruction box motif predicted to be recognized by the APC/C (Kitagawa et al., 2002; Fig. 4 A). We made a non-degradable IFY-1 by mutating two conserved residues in the destruction motif to alanine and fused it with GFP (GFP::IFY-1^DM^, Fig. 4 A). We generated transgenic GFP::IFY-1^DM^ lines and used methods we previously developed to grow and maintain *C. elegans* carrying toxic transgenes (Mitchell et al., 2014). As expected, overexpress3ion of GFP::IFY-1^DM^, but not GFP::IFY-1^WT^, caused embryonic lethality in multiple, independently generated strains (Fig. 4 B). Similar to previous studies in mouse oocytes (Herbert et al., 2003), GFP::IFY-1^DM^ causes obvious chromosome segregation and polar body extrusion defects (N= 28/31, Fig. 4 C). GFP::IFY-1^DM^ embryos shrink when dissected in high salt buffer indicating an eggshell permeability defect (N= 5/5), which is never observed in wildtype (N= 3/3, Fig. 4 D). Therefore, GFP::IFY-1^DM^ is dominant negative and causes phenotypes similar to inactivation of separase.

**Figure 4.**
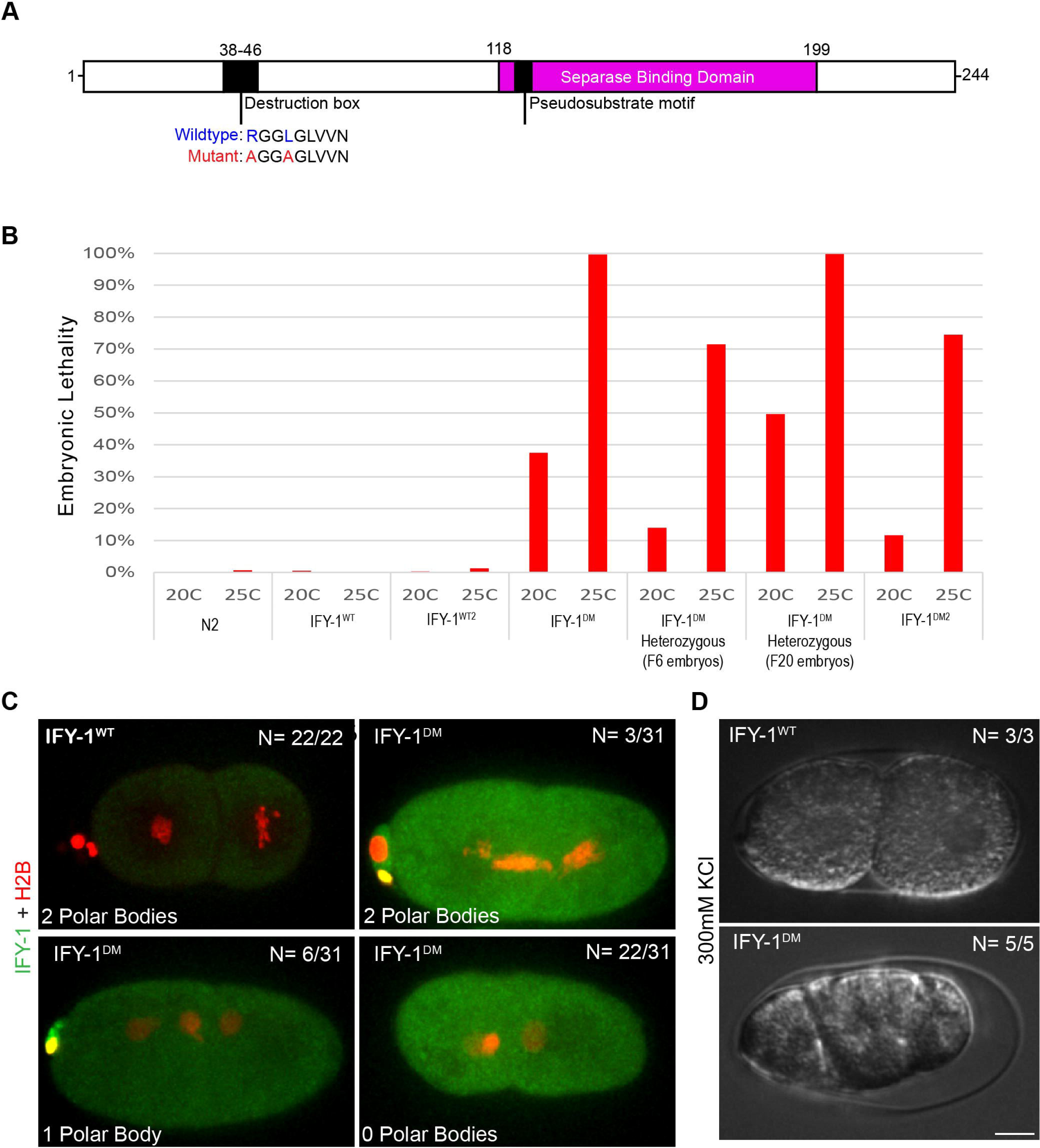
Generation and Characterization of GFP::IFY-1^DM^ in *C. elegans*. (A) Schematic of IFY-1 indicating the APC/C recognition motif (destruction box), within the unstructured region at the N-terminus. Separase binding domain indicates the region that was resolved in the structure of the separase/securin complex. Conserved residues in the D-box were mutated to alanine to prevent APC/C recognition. (B) Embryonic lethality of multiple independently generated transgenic IFY-1^WT^::GFP and IFY-1^DM^::GFP lines at 20°C and 25°C in homozygous or heterozygous animals. (C) In IFY-1^WT^::GFP, two-celled embryos always have two polar bodies. GFP::IFY-1^DM^ expression causes a spectrum of polar body extrusion defects with most embryos lacking any polar bodies, indicating defects in meiotic divisions. (D) IFY-1^WT^::GFP embryos that complete meiosis are not permeable, while IFY-1^DM^::GFP embryos all shrink in hyperosmotic solution, indicating permeability barrier defects. N = number of embryos scored. Scale bar: 10µm.

We examined GFP::IFY-1^DM^ localization during meiosis I with chromosome marker H2B::mCherry (Fig. 1 I-L). We observed that the GFP::IFY-1^DM^ localization pattern was identical to GFP::IFY-1^WT^ from prophase I through metaphase I, localizing to kinetochore cups and linear elements throughout the cortex (Fig. 1 E-G, I-K). As expected, GFP::IFY-1^DM^ was stable during anaphase I (Fig. 1 L, Fig. S 1 G) in contrast to the rapidly degraded GFP::IFY-1^WT^ (Fig. 1 H, M). GFP::IFY-1^DM^ began to accumulate on the spindle and remained strongly present on the spindle throughout anaphase I (Fig. 1 L). During anaphase, GFP::IFY-1^DM^ was not observed on vesicles, but remained prominent at the central spindle (Fig. 1 L). Therefore, GFP::IFY-1^DM^ behaves as expected for a mutant that specifically disrupts APC/C dependent degradation.

### IFY-1^DM^ causes chromosome segregation defects during anaphase I

Securin must be destroyed to allow chromosome segregation to occur. In many systems, expression of non-degradable securin causes cell division defects, (Cohen-Fix et al., 1996; Hagting et al., 2002; Herbert et al., 2003; Leismann et al., 2000; Zur & Brandeis, 2001), which has not been tested in worms. Therefore, we examined chromosome segregation during meiosis I in oocytes overexpressing GFP::IFY-1^WT^ or GFP::IFY-1^DM^ together with the chromosome marker H2B::mCherry (Fig. 5). In GFP::IFY-1^WT^ embryos, spindle rotation, which depends on APC/C activity (Crowder et al., 2015; Ellefson & McNally, 2011), occurs 68+/-7 seconds (N= 6) before anaphase onset. At anaphase onset, GFP::IFY-1^WT^ levels are rapidly decreasing and chromosomes move apart quickly (Fig 2 D, Fig. 5 A). In contrast, GFP::IFY-1^DM^ remains high on the spindle and accumulates at the midbivalent 56+/-6 seconds (N = 16) after spindle rotation. In stark contrast to WT, where reduced GFP::IFY-1^WT^ midbivalent localization can be seen approximately 5+/-2 seconds before chromosome movement (N= 6), chromosomes do not begin to move apart until approximately 115+/-8.5 seconds (N=23) after high levels of GFP::IFY-1^DM^ localize to the midbivalent (Fig. 2 E, Fig. 5 B). After this extended delay, chromosomes move poleward at a significantly slower rate than WT (Fig. 5 C). In addition, lagging chromosomes and severe chromosome segregation defects were observed in all movies of embryos expressing GFP::IFY-1^DM^ (N=18/18, Fig. 5 D), but similar defects were not observed in embryos expressing GFP::IFY-1^WT^ (N= 5/5, Fig. 5 D). Finally, in anaphase GFP::IFY-1^DM^ labels the central spindle in anaphase, which appears unstable and moves away from segregating chromosomes in late anaphase I (Fig. 5 B, t= 440 seconds), a phenotype not observed in WT. Therefore, GFP::IFY-1^DM^ causes severe chromosome segregation defects and a novel delay between the midbivalent localization of securin and the poleward movement of chromosomes.

**Figure 5.**
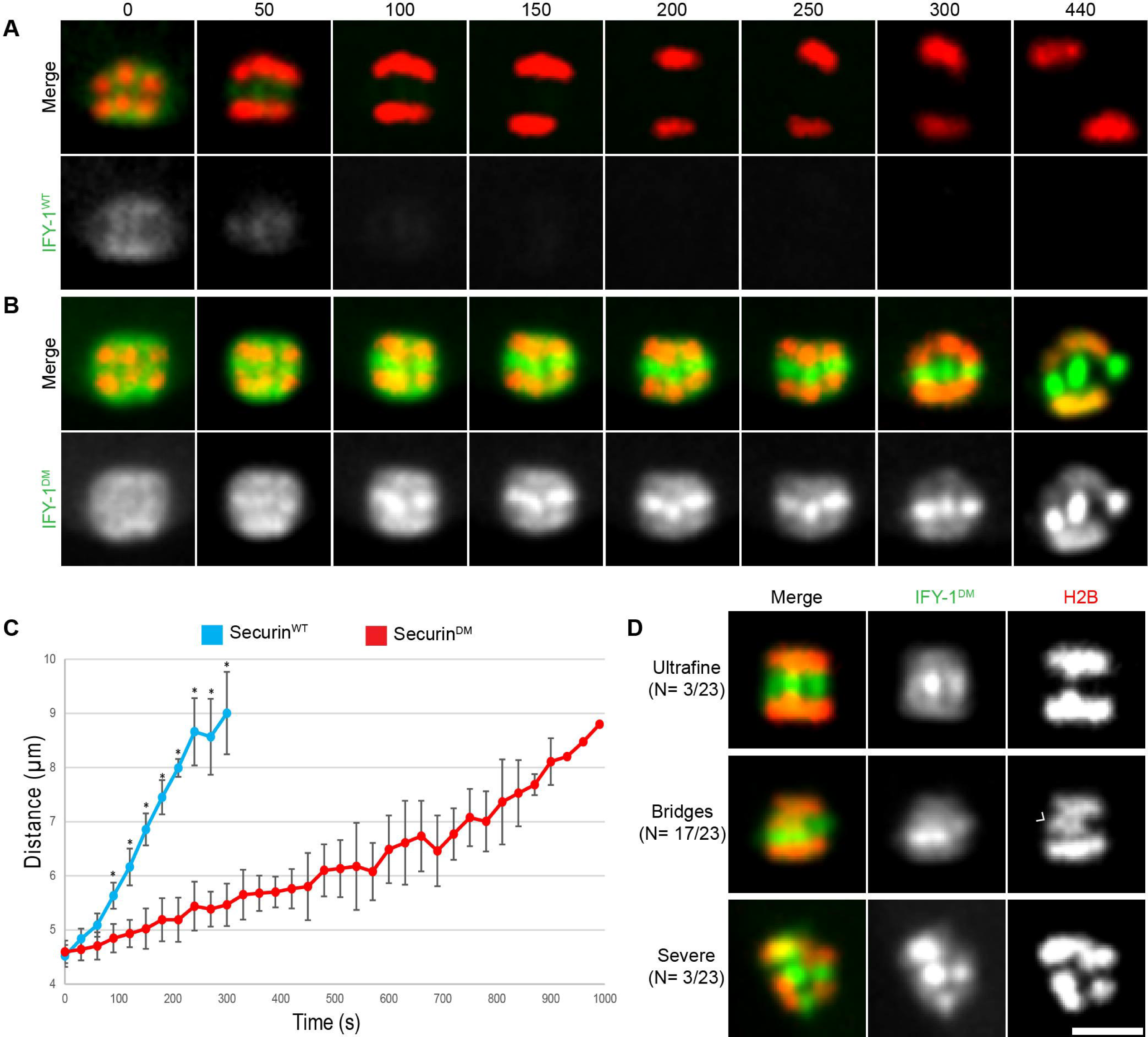
GFP::IFY-1^DM^ inhibits chromosome segregation during anaphase I. (A) IFY-1^WT^::GFP or (B) IFY-1^DM^::GFP (green) expressed together with H2B::mCherry (red) during meiosis I. Chromosomes move apart quickly in IFY-1^WT^::GFP, but are severely delayed in IFY-1^DM^::GFP embryos. Time (seconds) is relative to midbivalent localization. (C) Average distance between segregating chromosomes during anaphase I showing a significant delay in IFY-1^DM^::GFP embryos. The IFY-1^DM^::GFP curve starts after an extended delay of chromosome movement after midbivalent localization. N = 9 for IFY-1^WT^::GFP, N > 18 for IFY-1^DM^::GFP, asterisks denote a statistically significant difference. P-value = <0.05, error bars are standard deviations of the mean. (D) Frequency and representative images of ultrafine, chromosome bridging (caret) and severely impaired chromosome segregation defects observed in embryos expressing IFY-1^DM^::GFP (green, H2B::mCherry is red). N = number of embryos scored. Scale bar: 5µm.

### IFY-1^DM^ blocks cortical granule exocytosis during anaphase I

We next investigated whether securin degradation is required for separase to promote exocytosis, an important question for understanding the regulation of this process. Given that GFP::IFY-1^DM^ causes eggshell permeability (Fig. 4 D), we investigated whether it inhibited cortical granule exocytosis. We imaged embryos expressing GFP::IFY-1^WT^ or GFP::IFY-1^DM^ together with the cortical granule cargo protein, mCherry::CPG-2 during meiosis I (Fig. 6). In WT embryos, cortical granule exocytosis occurs within minutes of anaphase onset, depositing mCherry::CPG-2 into the eggshell (Fig. 6 A-C). In embryos expressing GFP::IFY-1^DM^, most of the mCherry::CPG-2 labeled cortical granules do not undergo exocytosis during anaphase I (Fig. 6 D-F). We quantified the number of vesicles released during anaphase, and found that GFP::IFY-1^DM^ embryos had a very high level of retained vesicles, which were completely released in GFP::IFY-1^WT^ embryos during anaphase I (Fig. 6 G). Therefore, GFP::IFY-1^DM^ expression causes a severe block of cortical granule exocytosis.

**Figure 6.**
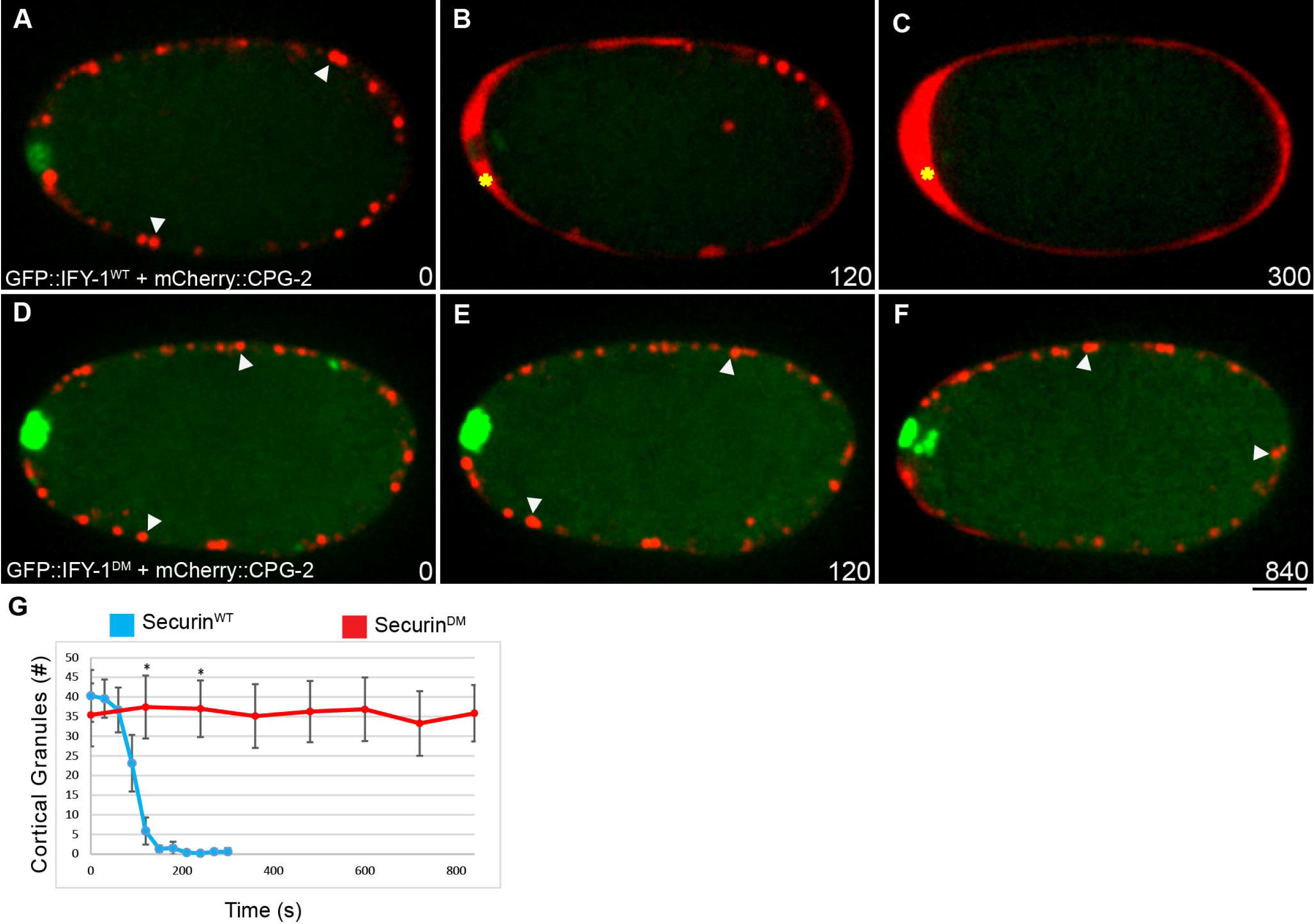
GFP::IFY-1^DM^ blocks cortical granule exocytosis in anaphase I. (A-C) IFY-1^WT^::GFP and (D-F) IFY-1^DM^::GFP (green) were co-expressed with the cortical granule cargo protein, mCherry::CPG-2 (red, white arrows). Time (seconds) is relative to midbivalent localization. In wildtype, the majority of cortical granules are exocytosed by 120 seconds after anaphase onset (B) and mCherry::CPG-2 is extracellular (yellow asterisk), and by the end of anaphase I (C) mCherry::CPG-2 is completely extracellular. In contrast, (E-F) in IFY-1^DM^::GFP embryos, cortical granules are not exocytosed even after several minutes (yellow asterisks indicate mCherry::CPG-2 labeled vesicles). (G) Quantification of cortical granules in a single spindle plane over time after anaphase onset. N = 6 for IFY-1^WT^::GFP, N = 7 for IFY-1^DM^::GFP. Asterisks denote a statistically significant difference, P-value < .0001. Error bars represent standard error of the mean. Scalebar: 10µm.

### IFY-1^DM^ inhibits SEP-1 vesicle localization in anaphase I

Our results demonstrate that securin is a potent inhibitor of separase function during chromosome segregation and exocytosis. While it is well established that stabilized securin blocks the protease activity of separase, we wanted to investigate whether it also affects separase localization. Therefore, we examined the localization dynamics of endogenously tagged SEP-1::mScarlet in oocytes and fertilized zygotes expressing GFP::IFY-1^WT^ or GFP::IFY^DM^ during meiosis I (Fig. 7). At early stages, from NEBD up until spindle rotation, SEP-1::mScarlet colocalizes normally with GFP::IFY-1^WT^ and GFP::IFY-1^DM^ on kinetochore cups and the spindle (Fig. 7 A, B). After spindle rotation and shortening, SEP-1::mScarlet colocalizes with GFP::IFY-1^WT^ and GFP::IFY-1^DM^ at spindle poles, but GFP::IFY-1^DM^ accumulates at higher levels (Fig. 7 A, B). Separase appears to normally localize to the midbivalent region and then persists on the anaphase spindle in the presence of either the rapidly lost GFP::IFY-1^WT^ or the highly accumulated GFP::IFY-1^DM^ (Fig. 7 A, B). Therefore, expression of GFP::IFY-1^DM^ severely inhibits chromosomes segregation without inhibiting the midbivalent or central spindle localization of separase.

**Figure 7.**
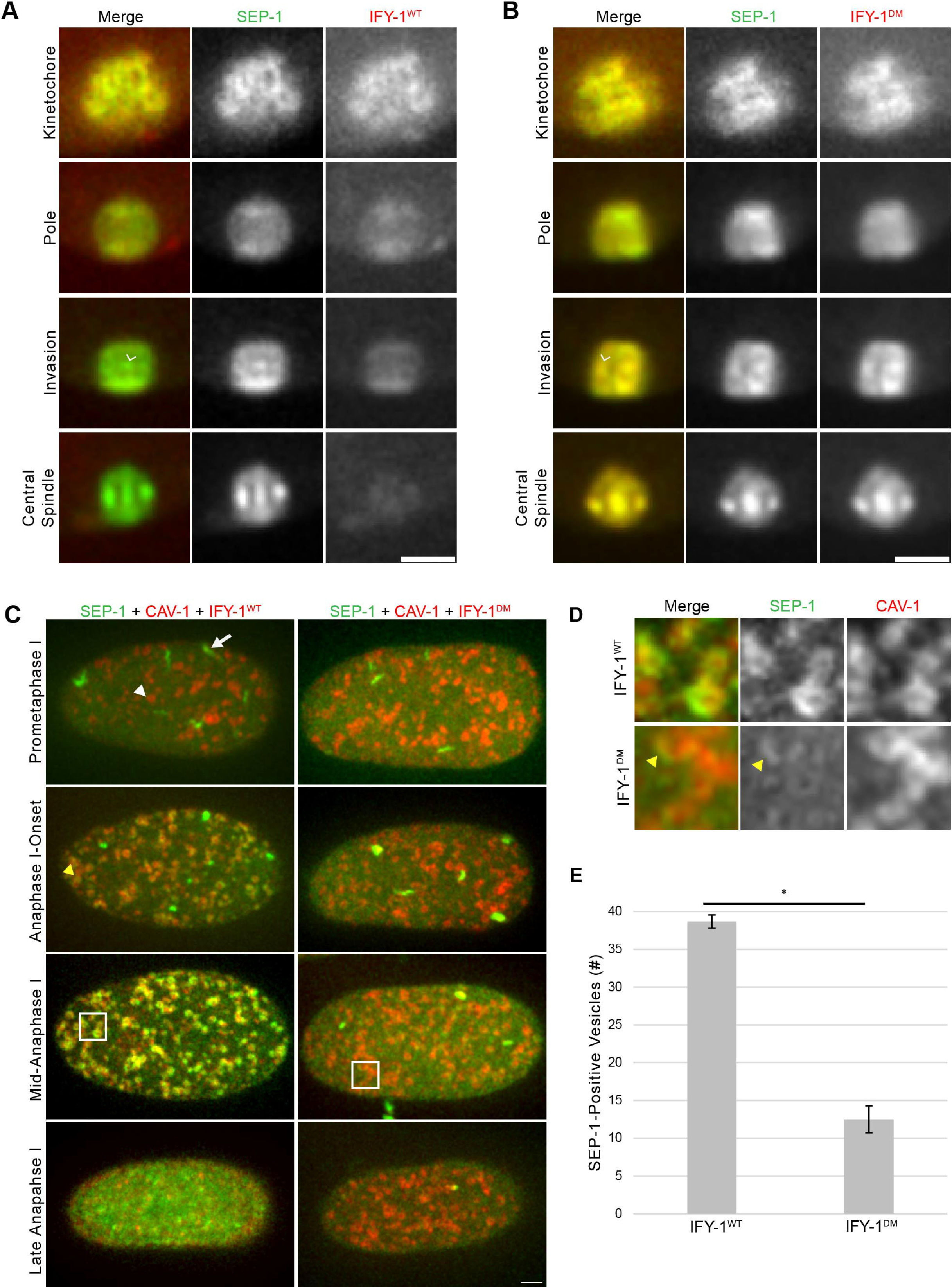
Separase localization to vesicles is reduced by GFP::IFY-1^DM^. (A) Representative spindle localization pattern of SEP-1::mScarlet (green) co-expressed with GFP::IFY-1^WT^ (red). Securin is rapidly degraded while separase moves from kinetochore cups to poles to the midbivalent (caret) at anaphase onset. (B) SEP-1::mScarlet colocalizes with stable GFP::IFY-1^DM^, in a similar pattern to wildtype at the midbivalent (arrowhead) and on the spindle in anaphase. (C) Max projections of cortical images of SEP-1::mScarlet (green) with the vesicle marker CAV-1::GFP (red) and either IFY-1^WT^::GFP or IFY-1^DM^::GFP (red). In prometaphase I SEP-1 localizes to linear elements (white arrow) but not vesicles (white arrowheads) in both conditions. At anaphase I onset, SEP-1::mScarlet begins to enrich on vesicles (yellow arrowhead) in GFP::IFY-1^WT^ but not GFP::IFY-1^DM^ embryos. By mid-anaphase I, SEP-1::mScarlet is fully enriched on vesicles in GFP::IFY-1^WT^ embryos. In GFP::IFY-1^DM^ embryos, SEP-1::mScarlet does not enrich on vesicles at anaphase I onset and only shows trace vesicle localization during anaphase I. (D) Magnified images of SEP-1::mScarlet vesicle localization from the 5µm^2^ regions indicated in (C) at mid-anaphase I in wildtype and mutant. SEP-1::mScarlet signal is reduced on vesicles when co-expressed with IFY-1^DM^::GFP, with only partial vesicle localization in rare cases (yellow arrowhead). (E) Quantification of vesicle-associated SEP-1::mScarlet signal in cortical planes at mid-anaphase I in GFP::IFY-1^WT^ (N = 11) and GFP::IFY-1^DM^ (N = 12) embryos. Asterisk denotes a statistically significant difference, P-value < .0001. Error bars represent standard error of the mean. Scale bar: 5µm.

Next, we investigated whether securin affects separase localization to vesicles. GFP::IFY-1^WT^ appears largely degraded by the time that separase localizes to vesicles, and we did not observe vesicle localization of GFP::IFY-1^DM^ (Fig. 1). Separase normally localizes to cortical granules immediately following anaphase I onset (Fig. 2 F). We imaged SEP-1::mScarlet in the presence of overexpressed GFP::IFY-1^WT^ or GFP::IFY-1^DM^ (Fig. 7). In GFP::IFY-1^WT^ embryos, SEP-1::mScarlet localized to vesicles normally (N=11/11 movies), but vesicles were not obviously observed in embryos expressing GFP::IFY-1^DM^ (N=11/12 movies). To more precisely examine separase localization to vesicles, we co-expressed the cortical granule marker CAV-1::GFP (Sato et al., 2008) together with GFP::IFY-1^WT^ or GFP::IFY-1^DM^ and examined SEP-1::mScarlet localization in the cortical-most planes of the embryo during anaphase I. During prometaphase I, SEP-1 localizes to linear elements in embryos expressing GFP::IFY-1^WT^ or GFP::IFY-1^DM^ (Fig. 7 C). After anaphase I onset, SEP-1::mScarlet accumulates on vesicles in GFP::IFY-1^WT^ but not GFP::IFY-1^DM^ embryos (Fig. 7 C, D). SEP-1::mScarlet showed prominent localization to CAV-1::GFP positive vesicles in mid anaphase when GFP::IFY-1^WT^ was expressed (Fig. 7 C, D). In contrast, SEP-1::mScarlet showed partial and reduced accumulation on a small subset of CAV-1::GFP positive vesicles in the presence of GFP::IFY-1^DM^ (Fig. 7 C, D). For WT, SEP-1::mScarlet rapidly accumulates on vesicles approximately 30 seconds after anaphase I onset, while 11/12 mutant embryos showed weak and partial vesicle localization after a significant delay (207 +/-37 seconds). We quantified the number of SEP-1::mScarlet positive vesicles in cortical planes in GFP::IFY-1^WT^ or GFP::IFY-1^DM^ expressing embryos (Fig. 7 E). In GFP::IFY-1^WT^ embryos, SEP-1::mScarlet localized to 40 +/-1 vesicles (N=11 embryos), while GFP::IFY-1^DM^ embryos had only 4 +/-1 SEP-1::mScarlet-positive vesicles (N=12 embryos). Therefore, GFP::IFY-1^DM^ interferes with the localization of SEP-1::mScarlet to cortical granules in anaphase I.

## Discussion

The proper regulation of separase activity is critical for the metaphase to anaphase transition. Previous studies have predominantly characterized separase regulation using *in vitro* biochemical assays to monitor its protease activity toward cohesin (Rosen et al., 2019). In addition to the activation of separase by APC/C mediated securin ubiquitination (Cohen-Fix et al., 1996; Funabiki et al., 1996), separase is regulated by autocleavage (Waizenegger et al., 2002), phosphorylation (Stemmann et al., 2001), prolyl isomerization (Hellmuth, Rata, et al., 2015), and CDK binding (Gorr et al., 2005) among others. However, the dynamic localization of separase and its regulation at different subcellular locations has not been well characterized. Underscoring the importance of separase localization, human cancer cells show aberrant separase nuclear localization (Meyer et al., 2009). Given that separase regulates centriole duplication (Tsou et al., 2009), anaphase spindle dynamics (Jensen et al., 2001) and vesicle exocytosis (Bembenek et al., 2007), different control mechanisms might be implemented in different ways to regulate these various processes. We have shown that separase is required for exocytosis of RAB-11 vesicles in meiosis and mitosis, which is important in both cases for successful cytokinesis in addition to eggshell formation during meiosis (Bembenek et al., 2007, 2010). Whether securin regulates cytokinesis has not been directly tested. A previous study suggested that expression of non-degradable securin causes chromosome segregation defects but does not affect cytokinesis (Zur & Brandeis, 2001). However, expression of non-cleavable cohesin causes chromosome segregation defects and cytokinesis failures (Hauf et al., 2001). This apparent discrepancy was partially addressed when a mechanism that prevents cytokinesis failures when chromatin bridges are present, called the abscission checkpoint, was identified (Mendoza et al., 2009). In this manuscript, we investigated whether and how securin might regulate vesicle exocytosis, which would directly contribute to cytokinesis events. Our results demonstrate that the spatiotemporal control of securin degradation and separase localization are critical for proper regulation of anaphase and directly regulate exocytosis.

To better understand separase regulation by securin, we defined the dynamics of the key events of the metaphase to anaphase transition with high spatiotemporal resolution. We demonstrate that although securin destruction begins a few minutes prior to anaphase onset in the cytoplasm, the population enriched on kinetochore structures in the spindle and in the cortex is rapidly lost shortly before anaphase onset. During the rapid securin degradation phase, separase undergoes a dynamic relocalization from kinetochore cups, to the spindle poles before enriching at the midbivalent where the specialized meiosis I cohesin complex resides (Severson & Meyer, 2014). Although separase had previously been shown to relocalize to the midbivalent region (Bembenek et al., 2010; Muscat et al., 2015), the timing of this event was not well characterized. We demonstrate that separase colocalizes with cohesin at the midbivalent seconds before cohesin is lost from chromosomes and the sister chromatids move poleward. Securin degradation is therefore completed to a sufficient extent to allow high separase activity when chromosomes come apart. The pole enriched localization of separase was not previously documented. During mitosis, separase was found to move from kinetochores to bulk chromatin during anaphase (Cabral et al., 2013), indicating that relocalization of separase may be important during meiosis and mitosis. We suspect that this dynamic relocalization reflects an active transport mechanism. Interestingly, inactivation of the spindle checkpoint involves dynein-mediated transport of checkpoint proteins from the kinetochore to the centrosome (Griffis et al., 2007; Howell et al., 2001). It will be interesting to determine whether this is related to the rapid relocalization of separase we document here. The mechanism required for the relocalization of separase to sites of action will be an important area of investigation for future studies.

The observation that separase and securin behave similarly on both kinetochore cups and linear elements in the cortex during prometaphase I is consistent with the well documented expectation that separase should be inactive during this time window. The kinetochore localization of separase and securin may reflect the local recruitment of separase to the vicinity of anaphase target sites while preventing it from directly interacting with substrates. The linear elements were first observed by staining for HIM-10 (Howe et al., 2001), and subsequently numerous outer kinetochore proteins were also found to localize to them (Hattersley et al., 2016; Monen et al., 2015; Pereira et al., 2018; Quiogue et al., 2023). Similar kinetochore linear element structures were observed in fly and bovine oocytes, suggesting these are widely conserved structures (Gilliland et al., 2007; Wu et al., 2023). Recent studies have shown that the outer kinetochore expands, forming a so-called fibrous corona, in order to facilitate microtubule capture at the early stages of mitosis (Kops & Gassmann, 2020). Many of the outer kinetochore proteins in the fibrous corona are also found in the cytoplasmic linear elements. Although a definitive function of linear elements remains elusive, they have recently been shown to regulate cortical microtubules to control plasma membrane dynamics during polar body extrusion (Quiogue et al., 2023). One interesting possibility is that the microtubule regulatory function of these cortical filaments could contribute to separase relocalization. Identifying kinetochore proteins that recruit separase to the chromosome and cortex and understanding the function of the linear elements will be major goals for future studies.

Our results reveal that the APC/C mediated destruction of securin not only activates separase proteolytically but also controls its spatial distribution in cells. Securin destruction in the cytoplasm begins early and likely reflects destruction of an excess pool of securin. Such a pool was found in mouse oocytes and was shown to regulate timing of anaphase onset (Thomas et al., 2021). Once the meiotic spindle begins to rotate and compact, which depends on the APC/C activity (Ellefson & McNally, 2011), kinetochore-localized securin is rapidly lost, and separase relocalizes to sites of action. Furthermore, inactivation of APC/C prevents separase from relocalizing to sites of action while securin depletion causes precocious relocalization. These results indicate that securin degradation by the APC/C is involved in timing the relocalization of separase at anaphase onset. APC/C regulates motor activity required for spindle translocation (Yang et al., 2005) and is essential for inactivation of cyclin dependent kinase (Pines, 2011). Therefore, APC/C may act on additional substrates besides securin to regulate separase relocalization. Consistent with this hypothesis, separase does not remain strictly associated with kinetochore cups and filaments when non-degradable securin is expressed in cells with APC/C activity, but relocalizes normally on the spindle while becoming diffuse in the cortex. Future studies will be needed to determine the targets of APC/C responsible for controlling separase localization.

It is well established that securin is a pseudo-substrate inhibitory chaperone of separase (Boland et al., 2017; Hellmuth et al., 2014; Hellmuth, Pöhlmann, et al., 2015; Hellmuth, Rata, et al., 2015; Hornig et al., 2002; Viadiu et al., 2005). As such it serves multiple regulatory roles including: (1) positively regulating the folding of separase (Hellmuth, Pöhlmann, et al., 2015); (2) enabling spindle and nuclear localization (Hornig et al., 2002); 3) binding to and inactivating the separase protease domain; and (4) acting as a competitive substrate of the APC/C to ensure proper timing of degradation and timing of events (Kamenz et al., 2015; Lu et al., 2014). We previously reported that both securin and separase RNAi inhibit cortical granule exocytosis (Bembenek et al., 2007). Given the multilayered function of securin, it was difficult to predict how its depletion might affect separase during exocytosis. A previous report indicated that separase failed to localize to cortical granules after securin depletion (Kimura & Kimura, 2012). However, we clearly document separase localization to vesicles when securin is depleted to various levels of depletion. Securin loss could cause separase to become unfolded and inactive, mislocalized, and/or prematurely active based on its known functions. The reduction in separase signal we observed after severe securin depletion could reflect mislocalization and/or unstable separase. In addition, we observe that securin depletion causes premature relocalization of separase to sites of action and causes premature loss of cohesin from chromosomes. Therefore, part of the securin phenotype is due to a premature activation of separase. It will be interesting to determine whether the failure of exocytosis when securin is depleted is partly due to premature cleavage of a vesicle substrate by separase.

To further characterize the role of securin in regulating the localization and activity of separase, we overexpressed non-degradable securin. Expression of GFP::IFY-1^DM^ should preserve the inhibitory chaperone function of securin and is not expected to affect the competitive substrate role since the APC/C recognition site is mutated. Therefore, GFP::IFY-1^DM^ specifically tests the effect of securin on separase in the presence of APC/C activity during anaphase. As expected, GFP::IFY-1^DM^ inhibits chromosome segregation and exocytosis, consistent with inhibiting the protease activity of separase. Importantly, we observe a novel chromosome segregation phenotype where separase localizes to the midbivalent but chromosomes do not move apart. Previously, this condition would not have been detected by only evaluating chromosome movement. Other mutant conditions that are expected to cause similar effects, for example an non-cleavable cohesin mutant, would be better analyzed by evaluating both separase localization and poleward chromosome movement. In our experiment, GFP::IFY-1^DM^ is expressed with endogenous securin also present in the cell. Our observation that GFP::IFY-1^DM^ accumulates to high levels on the spindle as anaphase progresses suggests that once endogenous securin is degraded, free GFP::IFY-1^DM^ might be capable of binding newly liberated separase and keeping it inactive, thus causing a severe phenotype. The finding that GFP::IFY-1^DM^ inhibits exocytosis is consistent with our previous work showing that protease dead separase inhibits cortical granule exocytosis (Bai & Bembenek, 2017). These results suggest that separase cleaves a substrate to promote exocytosis, which will be a major goal of future studies.

Interestingly, GFP::IFY-1^DM^ does not inhibit the midbivalent and spindle localization of separase, but significantly interferes with vesicle localization. We show that separase and securin colocalize with kinetochore proteins on chromosomes and in the cortex, suggesting that kinetochore proteins bind to the separase/securin complex. Since numerous kinetochore proteins are known to relocalize to the midbivalent and central spindle (Dumont et al., 2010), a complex of separase and GFP::IFY-1^DM^ may interact with kinetochore proteins at these sites in anaphase.

However, cortical filaments disappear during anaphase and kinetochore proteins are not known to localize to vesicles. Therefore, the failure of separase to enrich on cortical granules in the presence of GFP::IFY-1^DM^ suggests that securin blocks the domain of separase required for vesicle localization. Separase may bind directly to a substrate on vesicles, which would be inhibited by securin. We propose a model where separase is both kept inactive on kinetochores and out of reach of targeted substrates until the APC/C is activated. APC/C mediated securin degradation liberates separase protease activity and controls separase relocalization to sites of action at the spindle and cortex. On the spindle, active separase can interact with cohesin (and potentially other substrates) as well as kinetochore protein complexes. On vesicles, we propose that separase binds to a substrate and cleaves it to promote exocytosis. The spatiotemporal regulation of separase is an important facet of the metaphase to anaphase transition and may enable precise substrate cleavage by promoting a high local concentration of enzyme and substrate.

## Acknowledgements

CGC and Wormbase provided *C. elegans* strains and information, funded by the NIH (P40 OD010440) and NHGRI (U41 HG002223). We thank Julie Ahringer, Arshad Desai, Barth Grant, Tony Hyman, Karen Oegema, Martha Soto, and Asako Sugimoto for sharing strains and members of the Bembenek laboratory for comments and support. Funding was provided by the NIH grant R01 GM114471 to J.N.B.

## Video Legends

Video 1. **Chromosome dynamics at the metaphase-to-anaphase transition of meiosis I**. Single plane time lapse image series of embryos expressing H2B::mCherry with COH-3::GFP, SEP-1::GFP, GFP::IFY-1^WT^, or GFP::IFY-1^DM^. Time (seconds) is normalized to anaphase I onset (t = 0). Playback speed is 7 frames per second.

Video 2. **Separase dynamics at anaphase onset of meiosis I**. Single plane time lapse image series of embryos expressing SEP-1 (shown in green) with H2B, COH-3, or IFY-1^WT^ (shown in red). Time (seconds) is normalized to anaphase I onset (t = 0) as defined by SEP-1 midbivalent enrichment. Playback speed is 7 frames per second.

Video 3. **Securin and separase dynamics in the cortex during anaphase I**. Cortical time series from worms expressing SEP-1 (shown in green) with IFY-1 or RAB-11 (shown in red). Time (seconds) is normalized to anaphase I onset (t = 0), which occurs approximately 30 seconds before SEP-1 disappears from completely from kinetochore filaments. The SEP-1 + IFY-1 movie is a max projection acquired with a 60x objective, while SEP-1 + RAB-11 is a single plane movie acquired with a 100x objective. Playback speed is 7 frames per second.

Video 4. **GFP::IFY-1^DM^ blocks chromosome segregation during meiosis I**. Single plane time lapse image series of embryos expressing H2B::mCherry with GFP::IFY-1^WT^ or GFP::IFY-1^DM^. Time (seconds) is normalized to GFP enrichment at the midbivalent. Playback speed is 7 frames per second.

Video 5. **Separase localizes to the midbivalent and spindle when GFP::IFY-1^DM^ is overexpressed in anaphase I**. Single plane time lapse image series of embryos expressing SEP-1 (shown in green) with GFP::IFY-1^WT^ or GFP::IFY-1^DM^ (shown in red). Time is shown in seconds, t = 0 occurs when SEP-1 appears at the midbivalent. Playback speed is 5 frames per second.

Video 6. **GFP::IFY-1^DM^ prevents SEP-1 localization to cortical granules during anaphase I**. Cortical max projections from a time lapse image series of embryos expressing SEP-1 (green) and CAV-1 (red) with GFP::IFY-1^WT^ or GFP::IFY-1^DM^ (red). Time (seconds) is normalized to anaphase I onset (t = 0). Playback speed is 3 frames per second.

## Materials and Methods

### C. elegans Strains

Worm strains were maintained using standard protocols (Brenner, 1974; Mitchell et al., 2014). Some strains were obtained from the Caenorhabditis Genetics Center.

### Generation and Maintenance of GFP::IFY-1^DM^ Strains

The *ify-1* locus was PCR amplified from genomic DNA to include restriction sites at the 5’ (SpeI) and 3’end (MluI) for integration into the pJK3 plasmid, following established protocol (Gibson et al., 2009). The pJK3 plasmid allows N-terminal GFP fusion proteins expression through the *pie-1* promoter. The highly conserved arginine (Arg38) and leucine (Leu41) within the conserved destruction box were mutagenized to alanine (RxxL-> AxxA) using the Quickchange mutagenesis kit (Stratagene, La Jolla, CA). Worms were transformed with the pJK3 plasmid carrying *gfp::ify-1^dm^* using microparticle bombardment as previously described (Praitis et al., 2001). Multiple transformed worm lines were isolated and maintained using our protocol for the maintenance of worms harboring toxic transgenes (Mitchell et al., 2014).

Cloning and mutagenesis primers are listed below. Restriction sites are underlined and mutagenized residues are boldened.

*ify-1* Forward cloning prime with SpeI site: CGCTCTAGA<u>ACTAGT</u>ATGGAGGATCTAAAC

*ify-1* Reverse cloning prime with MluI: <u>ACGCGT</u>TCACAGGGGAAGGTTGGCTTCTTC

*ify-1^dm^*Forward destruction box mutagenesis primer: GGT**GCG**GGGCTGGTTGTAAACTCGTCA

*ify-1^dm^*Reverse destruction box mutagenesis primer: AGTCGAGTTTACAACCAGCCC**CGC**ACCGCC**AGC**AGAAGG

### Generation of endogenously tagged IFY-1^WT^::GFP and SEP-1::mScarlet

CRISPR/Cas9 was used to generate endogenously tag the wildtype *ify-1* locus at the N-terminus with GFP and the wildtype *sep-1* locus at the C-terminus with mScarlet, as previously described (Paix et al., 2015). The repair templates were amplified from the pDD282 and pMS050 plasmid (gifts from Bob Goldstein). The primer sequences and repair templates used are listed below. Underlined amino acids denote flexible linker sequences.

*ify-1::gfp* Forward: ACGACCTCCTCGCCGAAGAAGCCAACCTTCCCCTG<u>GGAGCATCGGGAGCC</u> GGAGCATCGGGAGCC

*ify-1::gfp* Reverse: AAACAGGTAGAAGAGGCTGACGTCGTGGGAAATCACTTGTAGAGCTCGTC CATTC

The *ify-1::gfp* guide RNA: GACGUCGUGGGAAAUCACAGGUUUUAGAGCUAUGCUGUUUUG

*sep-1::gfp* Forward: CAAGTGCCCGAACTCCATCAAGATCCCGAAATTTG<u>GGAGCATCGGGAGCC</u> <u>TCAGGAGCATCG</u>ATGGTCTCCAAGGG

*sep-1::gfp* Reverse: ACGATCCTTAAGATCCTTCGGGTCAGATTATATTACTTGTAGAGCTCGTCC ATTC

The *sep-1::gfp* guide RNA: CAGAUUAUAUUACAAAUUUCGUUUUAGAGCUAUGCUGUUUUG

### Creation of ySi12 *Ppie-1::GFP::coh-3*

The *coh-3* coding sequence and 3’ UTR were amplified from fosmid WRM068bC06 (Geneservice Ltd., Cambridge, UK) by PCR with primers AFS357 (GGGGACAGCTTTCTTGTACAAAGTGGctATGGTGATAAGCATCGATGTACC) and AFS358 (GGGGACAACTTTGTATAATAAAGTTGgcgcctttaaagctacctgtaac). The PCR product was cloned into pDONRP2R-P3 via a Gateway BP Cloning reaction (Thermo Fisher Scientific). A single missense mutation identified in the resulting plasmid and the parental fosmid was repaired by site-directed mutagenesis using primers AFS436 (/5Phos/TGAGTACTGAGAACTATGGTGTTTC) and AFS437 (/5Phos/AGTTGCTCGACTTCTTCgtac), yielding the error-free entry clone pAS139. pAS139 was used in a multi-site Gateway LR reaction together with entry plasmids pCG142 (*pie-1* intron*:pie-1* promoter in PDONRP4P1R) and pCM1.53 (GFP with worm codon bias and synthetic introns in pDONR201) (Addgene plasmids # 17246 and # 17250, gifts from Geraldine Seydoux) and destination vector pCFJ150 - pDESTttTi5605[R4-R3] (Addgene plasmid # 19329, gift from Erik Jorgensen). The resulting targeting vector, pAS142, was inserted into the *ttTi5605* Mos1 transposon site in strain EG4322 by MosSCI to create the single-copy integrated transgene *ySi12[Ppie-1::GFP::coh-3]* (Frøkjaer-Jensen et al., 2008; Merritt et al., 2008). *ySi12* encodes a functional GFP::COH-3 fusion, since it increases the embryonic viability of *coh-4(tm1857) coh-3(gk112)* double mutants from 3.1% (n=1246) to 91.8% (n=1522) and decreases male production from 36% (n=25) to 3.8% (n=1316). The low number of *coh-4 coh-3* animals scored for male production is due to the low rate of survival of these worms to adulthood; this phenotype is also rescued by the *ySi12* transgene.

### RNAi Treatments

Feeding RNAi was conducted as previously described using HT115 bacteria harboring the L4440 plasmid (Grishok et al., 2005). For *apc-2* and *ify-1* feeding RNAi, L4 hermaphrodites from WH416, JAB20, and JAB258 (Table 1) lines were plated onto the RNAi vector at 20°C or 25°C and phenotype severity and penetrance was assessed after 14-48hrs of treatment.

### Characterization of GFP::IFY-1^DM^ Lines

#### IFY-1 Degradation Curve and Spindle Curve

Degradation curve values are expressed as ratios reflecting the mean cytoplasmic GFP intensity in the newly fertilized oocyte relative to mean cytoplasmic GFP values in the −1 oocyte over time. Values for each timepoint correspond to an average of at least 5 independent movies for each IFY-1^WT^ or IFY-1^DM^ strain.

#### Embryonic Lethality

Lethality assays were performed as previously described (Mitchell et al., 2014). Lines expressing GFP::IFY-1^WT^ or GFP::IFY-1^DM^ were grown under identical conditions at 20L or 25L and embryo lethality was quantified. Lethality rates reflect the pooled average of embryonic lethality for each strain and condition after 24hrs.

#### Polar Body Extrusion Rate

The polar body extrusion assay was performed using embryos dissected from mothers homozygous for H2B::mCherry and GFP::IFY-1^WT^ or GFP::IFY-1^DM^ five generations removed from *gfp* RNAi feeding. We quantified two cell stage embryos to allow for the completion of meiosis and quantify polar bodies before the second polar body is internalized and degraded in older embryos (Fazeli et al., 2018).

#### Embryonic Osmotic Sensitivity

The osmotic sensitivity assay was performed by dissecting wildtype (N2) or homozygous GFP::IFY-1^DM^ mutant embryos in a hypertonic solution of 300mM KCl, as described (Knight et al., 2012). Animals were 5 generations removed from *gfp* RNAi feeding. Embryos were scored for normal appearance or obvious shrinkage.

### Live Cell Imaging

Live cell imaging data was collected using spinning disk confocal systems using either an inverted Nikon Eclipse microscope with a 60 X 1.40NA objective, a CSU-22 spinning disc system, and a Photometrics EM-CCD camera from Visitech International operated by MetaMorph software (Molecular Devices), or an inverted Nikon Eclipse Ti2-E with a 60 X 1.42NA objective and 100 X 1.45 NA objective, a CSU-X1 spinning disk system, and Andor iXon Life camera operated by NIS-Elements software (Nikon). Unless otherwise mentioned, live cell imaging was conducted at room temperature which was approximately 20°C. Image analysis and manipulation was performed in Fiji (National Institutes of Health), Adobe Photoshop and Adobe Illustrator (Adobe).

#### in Utero Live Cell Imaging

We used two immobilization methods to mount animals to image oocytes and embryos. The first method was an optimized nanoparticle-mediated immobilization technique based on a previously described protocol (Kim et al., 2013). This first strategy was used for Fig. 1 (A-C, E-H, I,J), Fig. 2 (A-D, H), Fig. S 1 (A, G), Fig. S 2 (B-E). We also used a chemical immobilization method by mounting worms in an M9 solution containing 5mM levamisole on a 2% agarose pad following standard protocol (Bai & Bembenek, 2017; Bembenek et al., 2007; Mitchell et al., 2014). We used this second strategy for documenting *control*, *apc-2*, and *ify-1* RNAi phenotypes, and for data presented in Fig. 2 (A-D”) and Fig. S 3 A.

#### ex utero Time Lapse Imaging of Meiosis I

Before eggshell formation, meiotic embryos are especially fragile to osmotic and mechanical perturbations (Stein & Golden, 2018). To minimize perturbations *ex utero*, meiotic embryos were dissected from hermaphrodites in blastomere culture media using the hanging drop mounting technique (Edgar & Goldstein, 2012). Parental carcasses were removed from the media along with bacteria to prevent toxic effects associated with their presence when left in the media (Bai & Bembenek, 2017; Bembenek et al., 2007; Mitchell et al., 2014). Cortical granules have weak autofluorescence under 488nm illumination that is quickly bleached after 25 exposures with standardized settings. Therefore, in conditions with weak GFP signal, we systematically performed an autofluorescence pre-bleach exposure before imaging GFP localization. For all *ex utero* imaging, L4 hermaphrodites were shifted from 20°C to 25°C for 18-24hrs before imaging at either room temperature or 25°C. This approach was used for acquiring data for Fig. 1 (D, M, N), Fig. 2 (E-I), Fig. 4 (C), Fig. 5 (A-D), Fig. 6 (A-G), Fig. (7A-D), Fig. S 1 (E,F), Fig. S 2 (A), and Movies 1-6.

### Quantifications

#### Fluorescence

Fluorescent values for degradation curves in Fig. 1 were acquired from single plane, *ex utero* movies of meiosis I embryos at approximately the same z-depth and using the same acquisition settings. The values represent the binned average signal from 2-5 independent movies, found in a 3-pixel diameter circle at the spindle, filament, and cytoplasm. The values for each movie and each timepoint at these regions of interest are averages of between 1-5 independent measurements per movie, minus the average background signal.

Fluorescent values for the securin degradation curves in Fig. S 2 were acquired from single plane, *in utero* movies of meiosis I. Worm age, imaging conditions (room temperature), and acquisition settings were identical for all data acquisition. Values represent the binned average of at least 2-4 independent movies, found in a 10-pixel diameter circle in the bulk cytoplasm. The values for each movie and each timepoint are averages of 3 independent cytoplasmic measurements, minus the average background signal.

Quantification of prometaphase I COH-3::GFP levels in Fig. S 2 were made from single plane, *in utero*, movies after RNAi treatments (*control*, *apc-2*, and *ify-1*). The calculated signal ratio corresponds to the midbivalent signal of the −1 oocyte to a prometaphase I oocyte in the spermatheca acquired in the same plane in a single image. A 3-pixel diameter circle region of COH-3::GFP signal was measured, subtracting average background signal within a 3-pixel diameter circle. Each condition is an average of 4-5 independent worms.

#### GFP::IFY-1^DM^ Reduction of Separase Vesicle Localization (*Fig. 7 D*)

We were unable to generate viable animals with homozygous GFP::IFY-1^DM^ and homozygous SEP-1::mScarlet, but heterozygous GFP::IFY-1^DM^ was viable when combined with heterozygous SEP-1::mScarlet. Extensive troubleshooting was taken to avoid phototoxicity and photobleaching, while simultaneously being able to evaluate SEP-1 vesicle localization. To time anaphase I onset, we imaged GFP alone using a single plane until GFP::IFY-1^DM^ was detected at the midbivalent. We then quickly switched to multiplane imaging (5µm steps x 3 planes) acquired every 45-120 seconds so that we could capture cortical planes over the course of an extended anaphase. Wildtype data was acquired only using multiplane *ex utero* imaging because embryos were not sensitive to phototoxic or photobleaching problems within the normal duration of anaphase I. Quantification of SEP-1::GFP signal at vesicle structures was performed on movies with z planes that had a similar circumference to ensure a similar region of cortex was quantified.

### Statistics

Calculations for p values were done in Microsoft Excel using Student’s t test (two-tailed, assuming unequal variance) to determine statistical significance.

---

**Figure S1.**
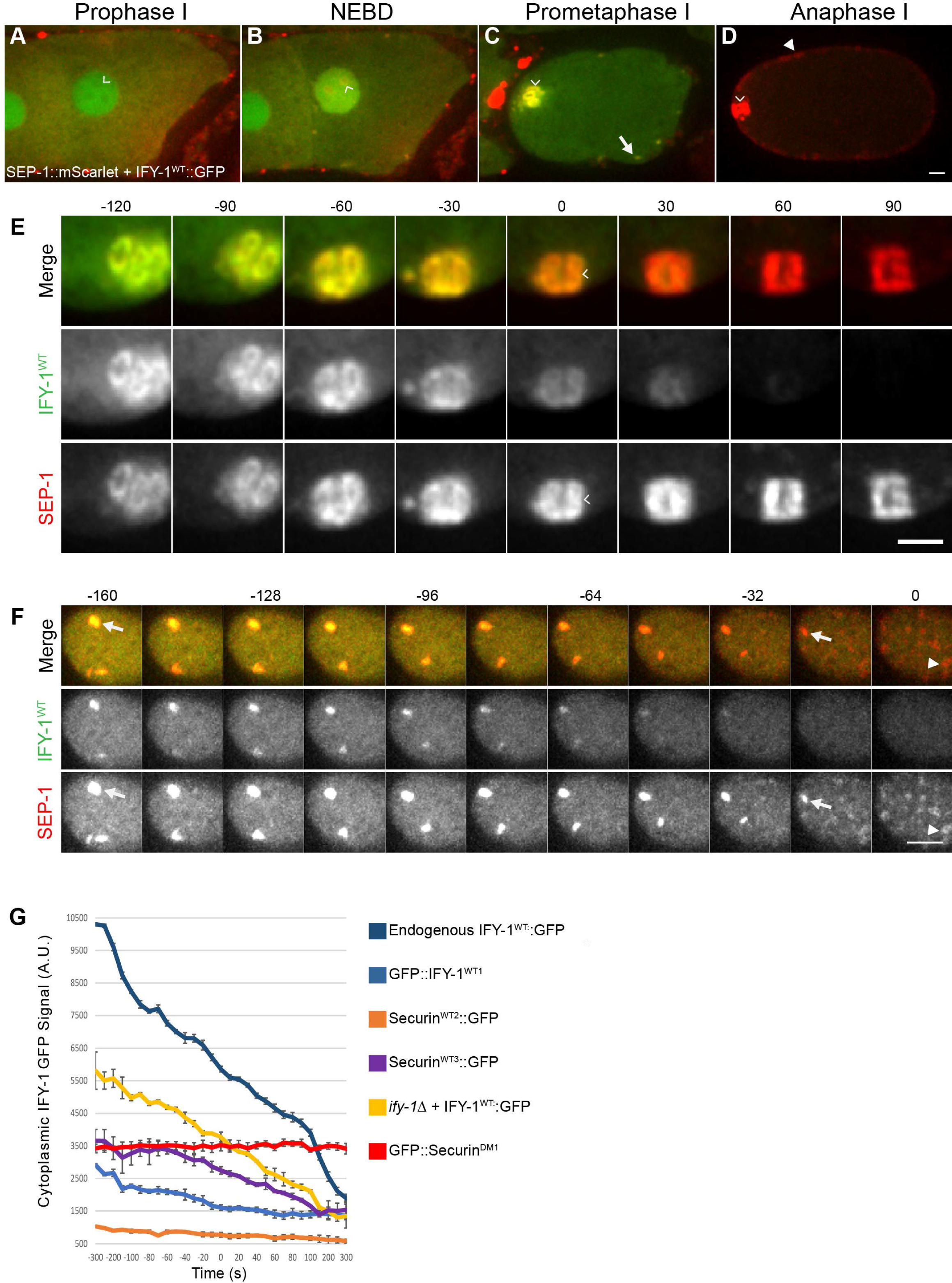
Dynamics of Separase and Securin during meiosis I. (**A-D**) Representative images of meiosis I in embryos co-expressing endogenously tagged SEP-1::mScarlet (red) and IFY-1::GFP (green). (A) At prophase I, IFY-1::GFP is present in both the nucleus (caret) and cytoplasm. SEP-1::mScarlet is cytoplasmic and excluded from the nucleus. (B) At NEBD, SEP-1::mScarlet enters the nucleus and colocalizes with IFY-1::GFP at kinetochores (caret). (C) In prometaphase I, SEP-1::mScarlet and IFY-1::GFP remain co-localized on kinetochore cups (caret), and linear elements (arrow) in the cortex. (D) During anaphase I, SEP-1 localizes to its sites of action at the central spindle (caret) and cortical granules (arrowhead) while IFY-1::GFP is mostly degraded. Scalebar: 10µm. (E) SEP-1::mScarlet (red) and IFY-1::GFP (green) colocalize on the spindle at the metaphase to anaphase transition. Securin is rapidly degraded while separase relocalizes to the midbivalent (caret) at anaphase onset (time shown in seconds, t = 0 is anaphase onset as defined by midbivalent accumulation) and remains highly enriched at the central spindle. (F) Cortical images of embryos expressing SEP-1::mScarlet (red) and IFY-1::GFP (green). SEP-1::mScarlet colocalizes with IFY-1::GFP on linear elements initially (arrows), but securin is largely degraded before separase relocalizes to vesicles (time shown in seconds, t = 0 marks prominent vesicle localization in mid anaphase). (G) Quantification of cytoplasmic securin levels in multiple WT securin GFP lines, compared with GFP::IFY-1^DM^. Regardless of expression levels, securin is rapidly degraded beginning prior to anaphase onset (t = 0 denotes chromosome segregation in WT and midbivalent localization in mutant). Scale bars: 5µm.

**Figure S2.**
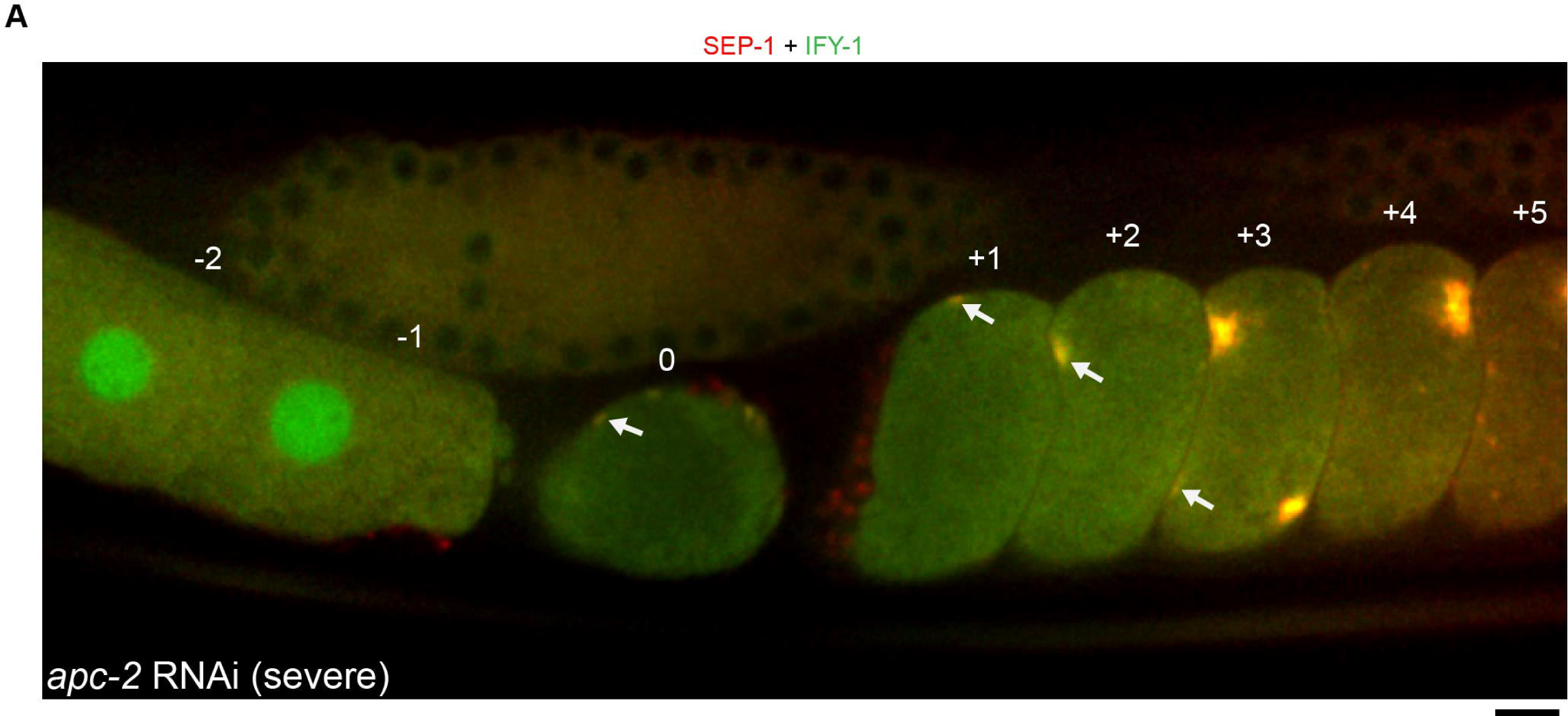
SEP-1 and IFY-1 persist on kinetochore structures after APC/C RNAi. (**A**) Representative image of embryos co-expressing endogenously tagged SEP-1::mScarlet (red) and endogenously tagged IFY-1::GFP (green) after *apc-2* RNAi. The image shown is a cortical slice from a z-series chosen to emphasize SEP-1 and IFY-1 persisting together on kinetochore filaments in the cortex (white arrows). Numbers correspond to relative position of oocytes and embryos in the hermaphrodite germline. Scale bar: 10µm.

**Figure S3.**
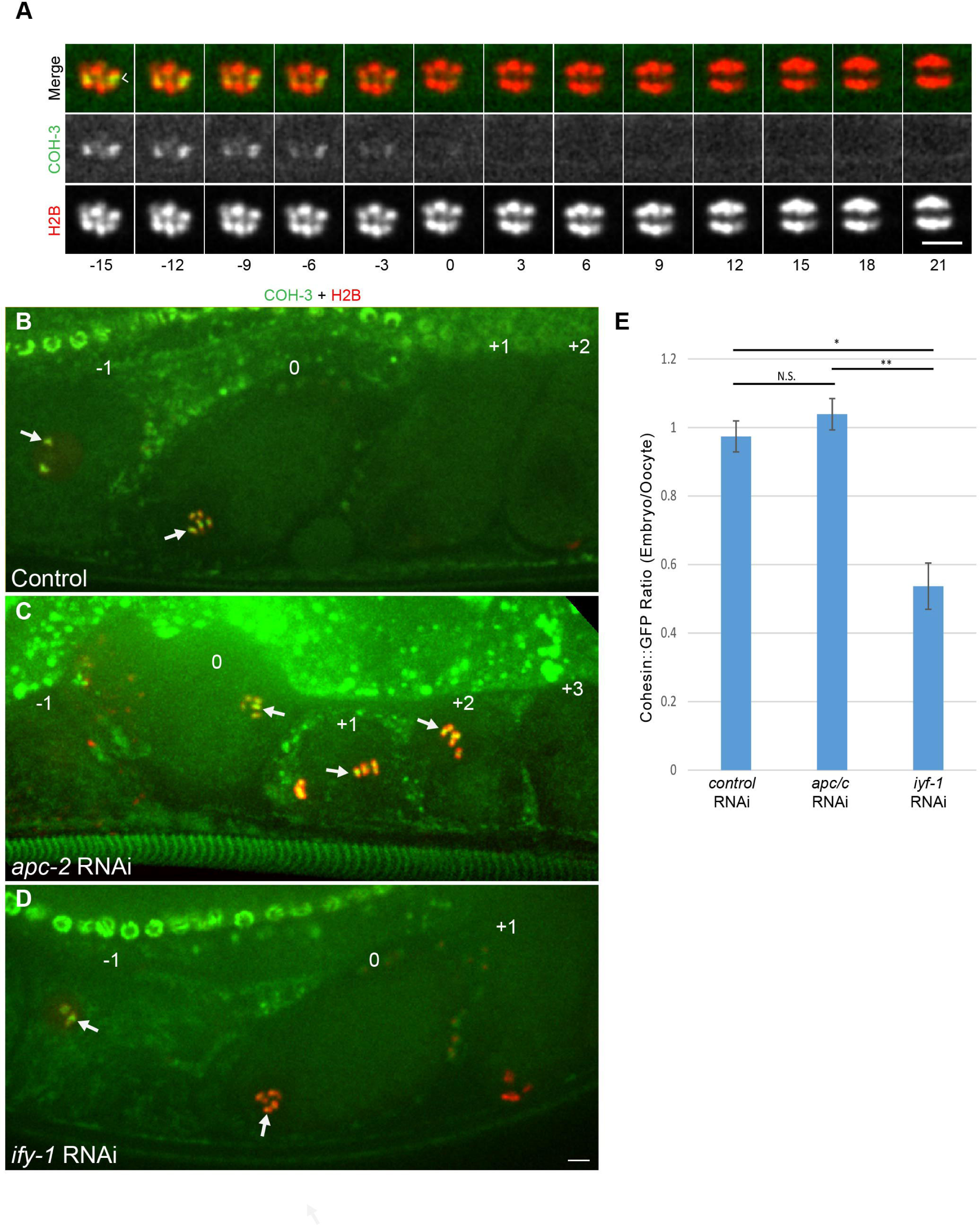
Meiosis I Cohesin dynamics and regulation during meiosis I. Representative images (A-D) of COH-3::GFP (green) with chromosome marker H2B::mCherry (red). (A) Kymograph from an *ex utero* time series at the metaphase-to-anaphase I transition. COH-3::GFP localizes to the midbivalent (caret) until shortly before anaphase I onset (t=0, time indicated in seconds). Representative images of the proximal germline after (B) control, (C) *apc-2* RNAi, and (D) *ify-1* RNAi. (B) In control, COH-3::GFP midbivalent signal is similar in oocytes (−1) and fertilized prometaphase I embryos in the spermatheca (labeled 0) (white arrows), but is absent from embryos after meiosis I in the uterus (+1, +2). (C) After *apc-2* RNAi, COH-3::GFP levels are retained at the midbivalent in prometaphase I embryos (labeled 0) and several arrested embryos in the uterus (+1, +2). (D) After *ify-1* RNAi, COH-3::GFP midbivalent signal prematurely reduced in prometaphase I embryos (labeled 0) relative to oocytes (−1) and is not observed in older embryos (+1). (E) Quantification of COH-3::GFP signal at the midbivalent in different conditions. The ratio between the COH-3::GFP signal on chromosomes in the prometaphase I embryo relative to the −1 oocyte in the same plane was calculated. The COH-3::GFP signal ratio was near 1 in control and *apc-2* RNAi (N=5, p>0.05) but was significantly reduced (about 50% reduction) after *ify-1* RNAi (N=4, p<0.05). Scale bars: 5µm.

**Table S1.**
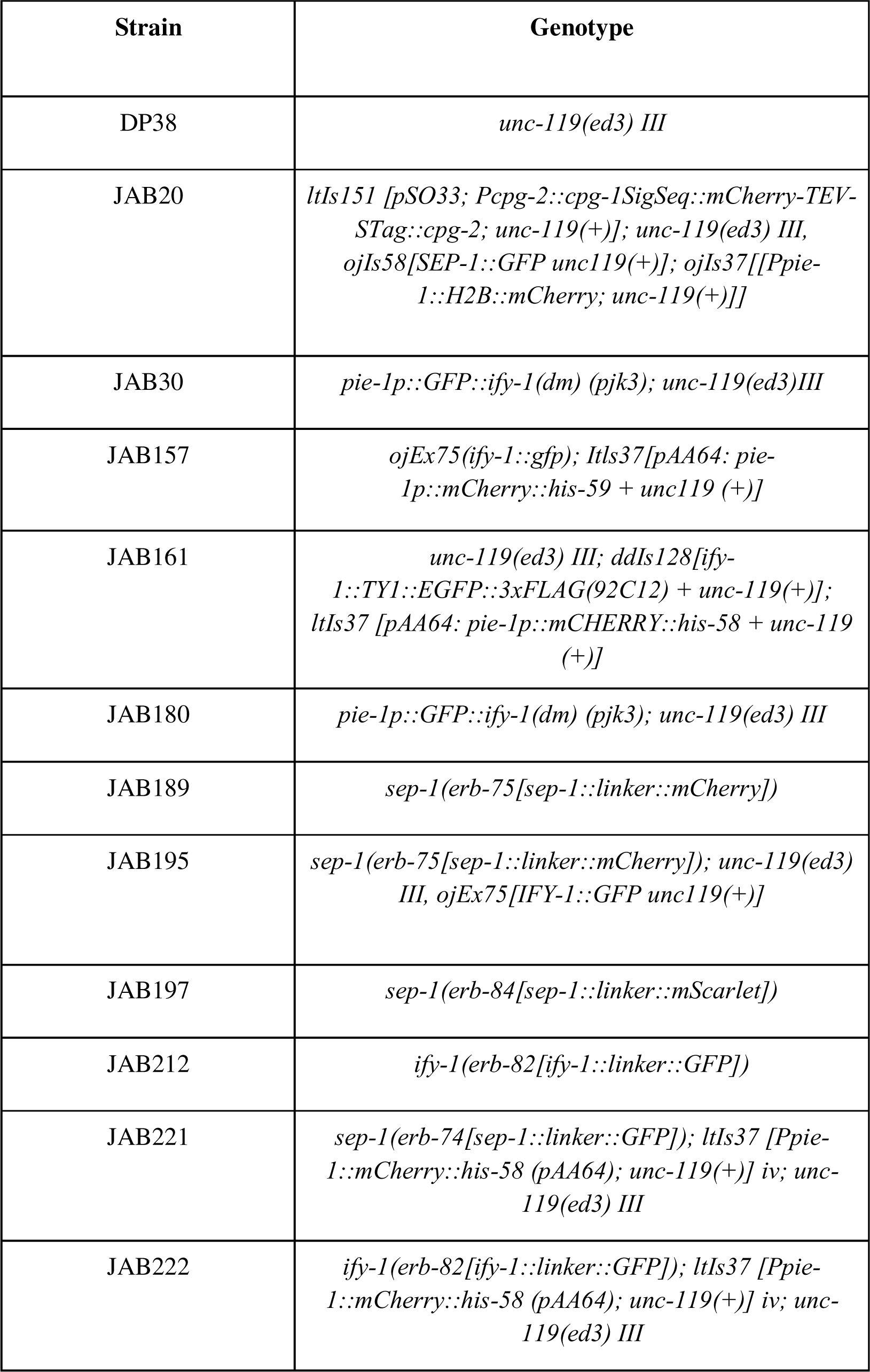
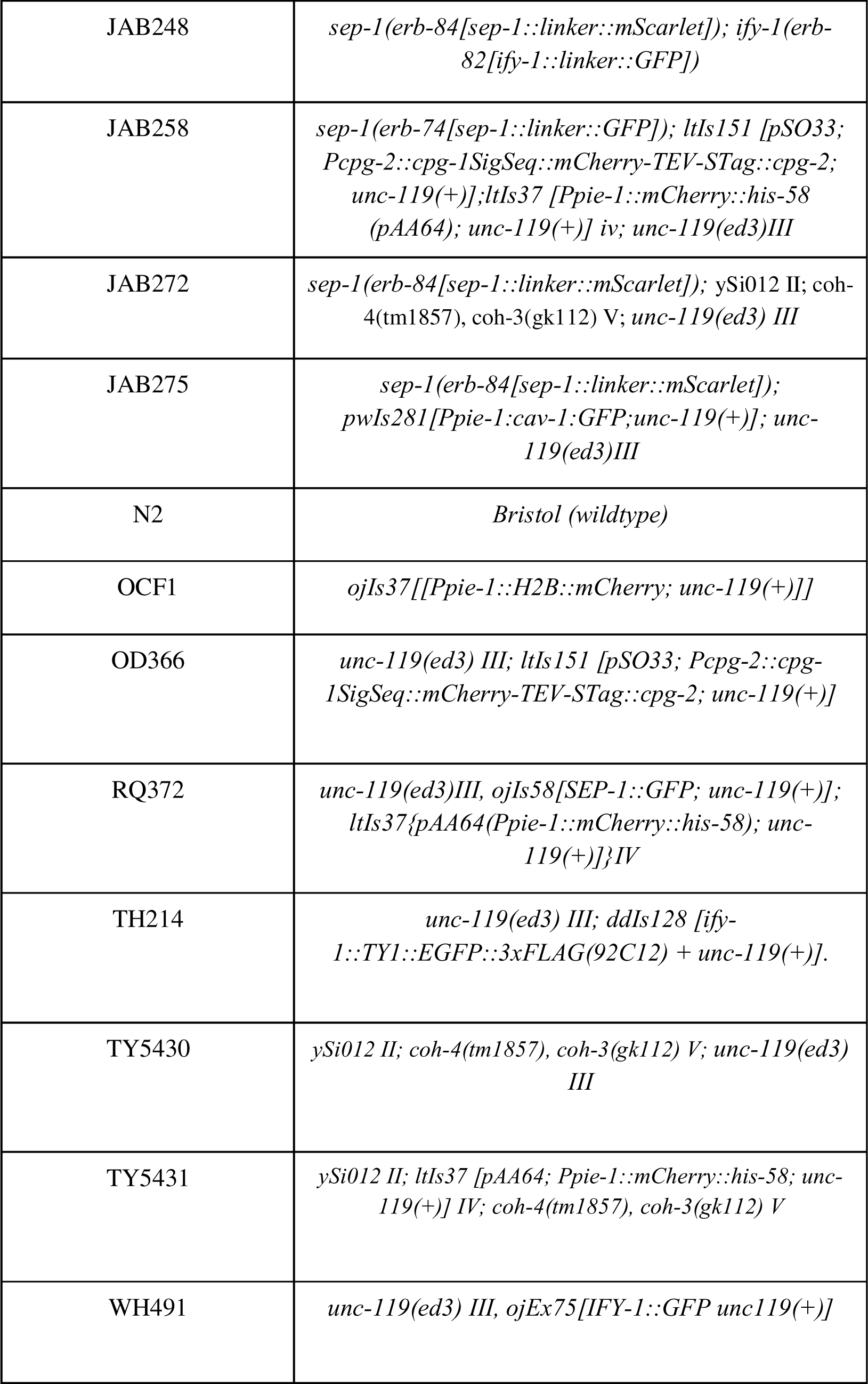
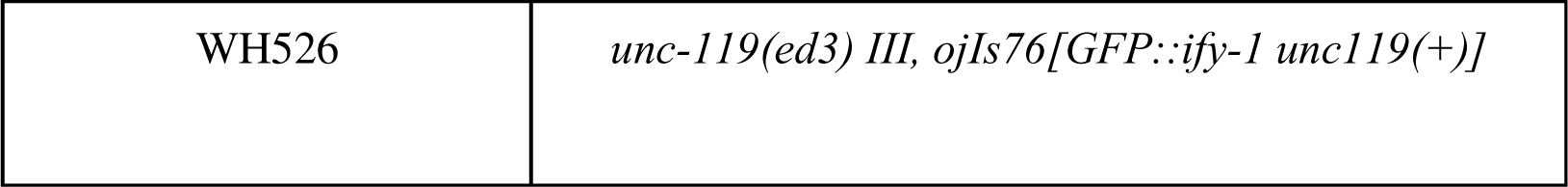
List of *C. elegans* strains used in this study.

## References

1. Agarwal, R., & Cohen-Fix, O. (2002). Phosphorylation of the mitotic regulator Pds1/securin by Cdc28 is required for efficient nuclear localization of Esp1/separase. Genes & Development, 16(11), 1371–1382. 10.1101/gad.971402

2. Agircan, F. G., & Schiebel, E. (2014). Sensors at Centrosomes Reveal Determinants of Local Separase Activity. PLOS Genetics, 10(10), e1004672. 10.1371/journal.pgen.1004672

3. Alfieri, C., Zhang, S., & Barford, D. (2017). Visualizing the complex functions and mechanisms of the anaphase promoting complex/cyclosome (APC/C). Open Biology, 7(11), 170204. 10.1098/rsob.170204

4. Bacac, M., Fusco, C., Planche, A., Santodomingo, J., Demaurex, N., Leemann-Zakaryan, R., Provero, P., & Stamenkovic, I. (2011). Securin and Separase Modulate Membrane Traffic by Affecting Endosomal Acidification. Traffic, 12(5), 615–626. 10.1111/j.1600-0854.2011.01169.x

5. Bai, X., & Bembenek, J. N. (2017). Protease dead separase inhibits chromosome segregation and RAB-11 vesicle trafficking. Cell Cycle, 16(20), 1902–1917. 10.1080/15384101.2017.1363936

6. Bembenek, J. N., Richie, C. T., Squirrell, J. M., Campbell, J. M., Eliceiri, K. W., Poteryaev, D., Spang, A., Golden, A., & White, J. G. (2007). Cortical granule exocytosis in C. elegans is regulated by cell cycle components including separase. Development (Cambridge, England), 134(21), 3837–3848. 10.1242/dev.011361

7. Bembenek, J. N., White, J. G., & Zheng, Y. (2010). A Role for Separase in the Regulation of RAB-11-positive Vesicles at the Cleavage Furrow and Midbody. Current BiologylJ: CB, 20(3), 259–264. 10.1016/j.cub.2009.12.045

8. Boland, A., Martin, T. G., Zhang, Z., Yang, J., Bai, X., Chang, L., Scheres, S. H. W., & Barford, D. (2017). Cryo-EM structure of a metazoan separase–securin complex at near-atomic resolution. Nature Structural & Molecular Biology, 24(4), Article 4. 10.1038/nsmb.3386

9. Brenner, S. (1974). The genetics of Caenorhabditis elegans. Genetics, 77(1), 71–94. 10.1093/genetics/77.1.71

10. Cabral, G., Sans, S. S., Cowan, C. R., & Dammermann, A. (2013). Multiple mechanisms contribute to centriole separation in C. elegans. Current Biology: CB, 23(14), 1380–1387. 10.1016/j.cub.2013.06.043

11. Chestukhin, A., Pfeffer, C., Milligan, S., DeCaprio, J. A., & Pellman, D. (2003). Processing, localization, and requirement of human separase for normal anaphase progression. Proceedings of the National Academy of Sciences of the United States of America, 100(8), Article 8. 10.1073/pnas.0730733100

12. Cohen-Fix, O., Peters, J. M., Kirschner, M. W., & Koshland, D. (1996). Anaphase initiation in Saccharomyces cerevisiae is controlled by the APC-dependent degradation of the anaphase inhibitor Pds1p. Genes & Development, 10(24), 3081– 3093. 10.1101/gad.10.24.3081

13. Crowder, M. E., Flynn, J. R., McNally, K. P., Cortes, D. B., Price, K. L., Kuehnert, P. A., Panzica, M. T., Andaya, A., Leary, J. A., & McNally, F. J. (2015). Dynactin-dependent cortical dynein and spherical spindle shape correlate temporally with meiotic spindle rotation in Caenorhabditis elegans. Molecular Biology of the Cell, 26(17), 3030–3046. 10.1091/mbc.E15-05-0290

14. Donangelo, I., Gutman, S., Horvath, E., Kovacs, K., Wawrowsky, K., Mount, M., & Melmed, S. (2006). Pituitary tumor transforming gene overexpression facilitates pituitary tumor development. Endocrinology, 147(10), 4781–4791. 10.1210/en.2006-0544

15. Dumont, J., Oegema, K., & Desai, A. (2010). A kinetochore-independent mechanism drives anaphase chromosome separation during acentrosomal meiosis. Nature Cell Biology, 12(9), Article 9. 10.1038/ncb2093

16. Edgar, L. G., & Goldstein, B. (2012). Culture and manipulation of embryonic cells. Methods in Cell Biology, 107, 151–175. 10.1016/B978-0-12-394620-1.00005-9

17. Ellefson, M. L., & McNally, F. J. (2011). CDK-1 inhibits meiotic spindle shortening and dynein-dependent spindle rotation in C. elegans. The Journal of Cell Biology, 193(7), 1229–1244. 10.1083/jcb.201104008

18. Fazeli, G., Stetter, M., Lisack, J. N., & Wehman, A. M. (2018). C. elegans Blastomeres Clear the Corpse of the Second Polar Body by LC3-Associated Phagocytosis. Cell Reports, 23(7), 2070–2082. 10.1016/j.celrep.2018.04.043

19. Foley, E. A., & Kapoor, T. M. (2013). Microtubule attachment and spindle assembly checkpoint signalling at the kinetochore. Nature Reviews Molecular Cell Biology, 14(1), Article 1. 10.1038/nrm3494

20. Frøkjaer-Jensen, C., Davis, M. W., Hopkins, C. E., Newman, B. J., Thummel, J. M., Olesen, S.-P., Grunnet, M., & Jorgensen, E. M. (2008). Single-copy insertion of transgenes in Caenorhabditis elegans. Nature Genetics, 40(11), 1375–1383. 10.1038/ng.248

21. Funabiki, H., Kumada, K., & Yanagida, M. (1996). Fission yeast Cut1 and Cut2 are essential for sister chromatid separation, concentrate along the metaphase spindle and form large complexes. The EMBO Journal, 15(23), 6617–6628.

22. Gibson, D. G., Young, L., Chuang, R.-Y., Venter, J. C., Hutchison, C. A., & Smith, H. O. (2009). Enzymatic assembly of DNA molecules up to several hundred kilobases. Nature Methods, 6(5), 343–345. 10.1038/nmeth.1318

23. Gilliland, W. D., Hughes, S. E., Cotitta, J. L., Takeo, S., Xiang, Y., & Hawley, R. S. (2007). The multiple roles of mps1 in Drosophila female meiosis. PLoS Genetics, 3(7), e113. 10.1371/journal.pgen.0030113

24. Gorbsky, G. J. (2015). The spindle checkpoint and chromosome segregation in meiosis. The FEBS Journal, 282(13), 2471–2487. 10.1111/febs.13166

25. Gorr, I. H., Boos, D., & Stemmann, O. (2005). Mutual inhibition of separase and Cdk1 by two-step complex formation. Molecular Cell, 19(1), Article 1. 10.1016/j.molcel.2005.05.022

26. Griffis, E. R., Stuurman, N., & Vale, R. D. (2007). Spindly, a novel protein essential for silencing the spindle assembly checkpoint, recruits dynein to the kinetochore. Journal of Cell Biology, 177(6), 1005–1015. 10.1083/jcb.200702062

27. Grishok, A. (2021). Small RNAs Worm Up Transgenerational Epigenetics Research. DNA, 1(2), 37–48. 10.3390/dna1020005

28. Grishok, A., Sinskey, J. L., & Sharp, P. A. (2005). Transcriptional silencing of a transgene by RNAi in the soma of C. elegans. Genes & Development, 19(6), 683–696. 10.1101/gad.1247705

29. Hagting, A., Den Elzen, N., Vodermaier, H. C., Waizenegger, I. C., Peters, J.-M., & Pines, J. (2002). Human securin proteolysis is controlled by the spindle checkpoint and reveals when the APC/C switches from activation by Cdc20 to Cdh1. Journal of Cell Biology, 157(7), 1125–1137. Scopus. 10.1083/jcb.200111001

30. Hattersley, N., Cheerambathur, D., Moyle, M., Stefanutti, M., Richardson, A., Lee, K.-Y., Dumont, J., Oegema, K., & Desai, A. (2016). A Nucleoporin Docks Protein Phosphatase 1 to Direct Meiotic Chromosome Segregation and Nuclear Assembly. Developmental Cell, 38(5), 463–477. 10.1016/j.devcel.2016.08.006

31. Hauf, S., Waizenegger, I. C., & Peters, J.-M. (2001). Cohesin Cleavage by Separase Required for Anaphase and Cytokinesis in Human Cells. Science, 293(5533), 1320– 1323. 10.1126/science.1061376

32. Heaney, A. P., Horwitz, G. A., Wang, Z., Singson, R., & Melmed, S. (1999). Early involvement of estrogen-induced pituitary tumor transforming gene and fibroblast growth factor expression in prolactinoma pathogenesis. Nature Medicine, 5(11), 1317–1321. 10.1038/15275

33. Hellmuth, S., Böttger, F., Pan, C., Mann, M., & Stemmann, O. (2014). PP2A delays APC/C-dependent degradation of separase-associated but not free securin. The EMBO Journal, 33(10), Article 10. 10.1002/embj.201488098

34. Hellmuth, S., Pöhlmann, C., Brown, A., Böttger, F., Sprinzl, M., & Stemmann, O. (2015). Positive and negative regulation of vertebrate separase by Cdk1-cyclin B1 may explain why securin is dispensable. The Journal of Biological Chemistry, 290(12), Article 12. c

35. Hellmuth, S., Rata, S., Brown, A., Heidmann, S., Novak, B., & Stemmann, O. (2015). Human chromosome segregation involves multi-layered regulation of separase by the peptidyl-prolyl-isomerase Pin1. Molecular Cell, 58(3), Article 3. 10.1016/j.molcel.2015.03.025

36. Herbert, M., Levasseur, M., Homer, H., Yallop, K., Murdoch, A., & McDougall, A. (2003). Homologue disjunction in mouse oocytes requires proteolysis of securin and cyclin B1. Nature Cell Biology, 5(11), Article 11. 10.1038/ncb1062

37. Horner, V. L., & Wolfner, M. F. (2008). Transitioning from egg to embryo: Triggers and mechanisms of egg activation. Developmental Dynamics, 237(3), 527–544. 10.1002/dvdy.21454

38. Hornig, N. C. D., Knowles, P. P., McDonald, N. Q., & Uhlmann, F. (2002). The Dual Mechanism of Separase Regulation by Securin. Current Biology, 12(12), Article 12. 10.1016/S0960-9822(02)00847-3

39. Howe, M., McDonald, K. L., Albertson, D. G., & Meyer, B. J. (2001). Him-10 Is Required for Kinetochore Structure and Function on Caenorhabditis elegans Holocentric Chromosomes. Journal of Cell Biology, 153(6), 1227–1238. 10.1083/jcb.153.6.1227

40. Howell, B. J., McEwen, B. F., Canman, J. C., Hoffman, D. B., Farrar, E. M., Rieder, C. L., & Salmon, E. D. (2001). Cytoplasmic dynein/dynactin drives kinetochore protein transport to the spindle poles and has a role in mitotic spindle checkpoint inactivation. Journal of Cell Biology, 155(7), 1159–1172. 10.1083/jcb.200105093

41. Jensen, S., Segal, M., Clarke, D. J., & Reed, S. I. (2001). A novel role of the budding yeast separin Esp1 in anaphase spindle elongation: Evidence that proper spindle association of Esp1 is regulated by Pds1. The Journal of Cell Biology, 152(1), 27–40. 10.1083/jcb.152.1.27

42. Kamenz, J., Mihaljev, T., Kubis, A., Legewie, S., & Hauf, S. (2015). Robust Ordering of Anaphase Events by Adaptive Thresholds and Competing Degradation Pathways. Molecular Cell, 60(3), Article 3. 10.1016/j.molcel.2015.09.022

43. Kim, E., Sun, L., Gabel, C. V., & Fang-Yen, C. (2013). Long-Term Imaging of Caenorhabditis elegans Using Nanoparticle-Mediated Immobilization. PLOS ONE, 8(1), e53419. 10.1371/journal.pone.0053419

44. Kimura, K., & Kimura, A. (2012). Rab6 is required for the exocytosis of cortical granules and the recruitment of separase to the granules during the oocyte-to-embryo transition in Caenorhabditis elegans. Journal of Cell Science, 125(Pt 23), 5897–5905. 10.1242/jcs.116400

45. King, R. W., Glotzer, M., & Kirschner, M. W. (1996). Mutagenic analysis of the destruction signal of mitotic cyclins and structural characterization of ubiquitinated intermediates. Molecular Biology of the Cell, 7(9), 1343–1357. 10.1091/mbc.7.9.1343

46. Kitagawa, R., Law, E., Tang, L., & Rose, A. M. (2002). The Cdc20 homolog, FZY-1, and its interacting protein, IFY-1, are required for proper chromosome segregation in Caenorhabditis elegans. Current Biology: CB, 12(24), 2118–2123. 10.1016/s0960-9822(02)01392-1

47. Knight, A. J., Johnson, N. M., & Behm, C. A. (2012). VHA-19 Is Essential in Caenorhabditis elegans Oocytes for Embryogenesis and Is Involved in Trafficking in Oocytes. PLOS ONE, 7(7), e40317. 10.1371/journal.pone.0040317

48. Kops, G. J. P. L., & Gassmann, R. (2020). Crowning the Kinetochore: The Fibrous Corona in Chromosome Segregation. Trends in Cell Biology, 30(8), 653–667. 10.1016/j.tcb.2020.04.006

49. Lara-Gonzalez, P., Pines, J., & Desai, A. (2021). Spindle assembly checkpoint activation and silencing at kinetochores. Seminars in Cell & Developmental Biology, 117, 86–98. 10.1016/j.semcdb.2021.06.009

50. Leismann, O., Herzig, A., Heidmann, S., & Lehner, C. F. (2000). Degradation of Drosophila PIM regulates sister chromatid separation during mitosis. Genes & Development, 14(17), 2192–2205. 10.1101/gad.176700

51. Liu, M. (2011). The biology and dynamics of mammalian cortical granules. Reproductive Biology and Endocrinology, 9(1), 149. 10.1186/1477-7827-9-149

52. Liu, M., Sims, D., Calarco, P., & Talbot, P. (2003). Biochemical heterogeneity, migration, and pre-fertilization release of mouse oocyte cortical granules. Reproductive Biology and Endocrinology: RB&E, 1, 77. 10.1186/1477-7827-1-77

53. Lu, D., Hsiao, J. Y., Davey, N. E., Van Voorhis, V. A., Foster, S. A., Tang, C., & Morgan, D. O. (2014). Multiple mechanisms determine the order of APC/C substrate degradation in mitosis. The Journal of Cell Biology, 207(1), 23–39. 10.1083/jcb.201402041

54. Lui, D. Y., & Colaiácovo, M. P. (2013). Meiotic Development in Caenorhabditis elegans. Advances in Experimental Medicine and Biology, 757, 133–170. 10.1007/978-1-4614-4015-4_6

55. MacKenzie, A., Vicory, V., & Lacefield, S. (2023). Meiotic cells escape prolonged spindle checkpoint activity through kinetochore silencing and slippage. PLOS Genetics, 19(4), e1010707. 10.1371/journal.pgen.1010707

56. McAinsh, A. D., & Kops, G. J. P. L. (2023). Principles and dynamics of spindle assembly checkpoint signalling. Nature Reviews Molecular Cell Biology, 24(8), Article 8. 10.1038/s41580-023-00593-z

57. McCarter, J., Bartlett, B., Dang, T., & Schedl, T. (1999). On the control of oocyte meiotic maturation and ovulation in Caenorhabditis elegans. Developmental Biology, 205(1), 111–128.o

58. McNally, K. L., & McNally, F. J. (2005). Fertilization initiates the transition from anaphase I to metaphase II during female meiosis in C. elegans. Developmental Biology, 282(1), 218–230. 10.1016/j.ydbio.2005.03.009

59. Mello, C. C., Schubert, C., Draper, B., Zhang, W., Lobel, R., & Priess, J. R. (1996). The PIE-1 protein and germline specification in C. elegans embryos. Nature, 382(6593), 710–712. 10.1038/382710a0

60. Mendoza, M., Norden, C., Durrer, K., Rauter, H., Uhlmann, F., & Barral, Y. (2009). A mechanism for chromosome segregation sensing by the NoCut checkpoint. Nature Cell Biology, 11(4), 477–483. 10.1038/ncb1855

61. Merritt, C., Rasoloson, D., Ko, D., & Seydoux, G. (2008). 3’ UTRs are the primary regulators of gene expression in the C. elegans germline. Current Biology: CB, 18(19), 1476–1482. 10.1016/j.cub.2008.08.013

62. Meyer, R., Fofanov, V., Panigrahi, A. K., Merchant, F., Zhang, N., & Pati, D. (2009). Overexpression and Mislocalization of the Chromosomal Segregation Protein Separase in Multiple Human Cancers. Clinical Cancer ResearchlJ: An Official Journal of the American Association for Cancer Research, 15(8), 2703–2710. 10.1158/1078-0432.CCR-08-2454

63. Mitchell, D. M., Uehlein-Klebanow, L. R., & Bembenek, J. N. (2014). Protease-Dead Separase Is Dominant Negative in the C. elegans Embryo. PLOS ONE, 9(9), e108188. 10.1371/journal.pone.0108188

64. Mogessie, B., Scheffler, K., & Schuh, M. (2018). Assembly and Positioning of the Oocyte Meiotic Spindle. Annual Review of Cell and Developmental Biology, 34(1), 381–403. 10.1146/annurev-cellbio-100616-060553

65. Monen, J., Hattersley, N., Muroyama, A., Stevens, D., Oegema, K., & Desai, A. (2015). Separase Cleaves the N-Tail of the CENP-A Related Protein CPAR-1 at the Meiosis I Metaphase-Anaphase Transition in C. elegans. PloS One, 10(4), e0125382. 10.1371/journal.pone.0125382

66. Moschou, P. N., Smertenko, A. P., Minina, E. A., Fukada, K., Savenkov, E. I., Robert, S., Hussey, P. J., & Bozhkov, P. V. (2013). The caspase-related protease separase (extra spindle poles) regulates cell polarity and cytokinesis in Arabidopsis. The Plant Cell, 25(6), 2171–2186. 10.1105/tpc.113.113043

67. Musacchio, A. (2015). The Molecular Biology of Spindle Assembly Checkpoint Signaling Dynamics. Current Biology: CB, 25(20), R1002–1018. 10.1016/j.cub.2015.08.051

68. Muscat, C. C., Torre-Santiago, K. M., Tran, M. V., Powers, J. A., & Wignall, S. M. (2015). Kinetochore-independent chromosome segregation driven by lateral microtubule bundles. eLife, 4, e06462. 10.7554/eLife.06462

69. Nam, H.-J., & van Deursen, J. M. (2014). Cyclin B2 and p53 control proper timing of centrosome separation. Nature Cell Biology, 16(6), Article 6. 10.1038/ncb2952

70. Olson, S. K., Greenan, G., Desai, A., Müller-Reichert, T., & Oegema, K. (2012). Hierarchical assembly of the eggshell and permeability barrier in C. elegans. The Journal of Cell Biology, 198(4), 731–748. 10.1083/jcb.201206008

71. Paix, A., Folkmann, A., Rasoloson, D., & Seydoux, G. (2015). High Efficiency, Homology-Directed Genome Editing in Caenorhabditis elegans Using CRISPR-Cas9 Ribonucleoprotein Complexes. Genetics, 201(1), 47–54. 10.1534/genetics.115.179382

72. Pereira, C., Reis, R. M., Gama, J. B., Celestino, R., Cheerambathur, D. K., Carvalho, A. X., & Gassmann, R. (2018). Self-Assembly of the RZZ Complex into Filaments Drives Kinetochore Expansion in the Absence of Microtubule Attachment. Current Biology: CB, 28(21), Article 21. 10.1016/j.cub.2018.08.056

73. Pines, J. (2011). Cubism and the cell cycle: The many faces of the APC/C. Nature Reviews. Molecular Cell Biology, 12(7), 427–438. 10.1038/nrm3132

74. Praitis, V., Casey, E., Collar, D., & Austin, J. (2001). Creation of Low-Copy Integrated Transgenic Lines in Caenorhabditis elegans. Genetics, 157(3), 1217–1226. 10.1093/genetics/157.3.1217

75. Quarato, P., Singh, M., Bourdon, L., & Cecere, G. (2022). Inheritance and maintenance of small RNA-mediated epigenetic effects. BioEssays: News and Reviews in Molecular, Cellular and Developmental Biology, 44(6), e2100284. 10.1002/bies.202100284

76. Quiogue, A. R., Sumiyoshi, E., Fries, A., Chuang, C.-H., & Bowerman, B. (2023). Microtubules oppose cortical actomyosin-driven membrane ingression during C. elegans meiosis I polar body extrusion. PLOS Genetics, 19(10), e1010984. 10.1371/journal.pgen.1010984

77. Richie, C. T., Bembenek, J. N., Chestnut, B., Furuta, T., Schumacher, J. M., Wallenfang, M., & Golden, A. (2011). Protein phosphatase 5 is a negative regulator of separase function during cortical granule exocytosis in C. elegans. Journal of Cell Science, 124(Pt 17), 2903–2913. 10.1242/jcs.073379

78. Robker, R. L., Hennebold, J. D., & Russell, D. L. (2018). Coordination of Ovulation and Oocyte Maturation: A Good Egg at the Right Time. Endocrinology, 159(9), 3209–3218. 10.1210/en.2018-00485

79. Rosen, L. E., Klebba, J. E., Asfaha, J. B., Ghent, C. M., Campbell, M. G., Cheng, Y., & Morgan, D. O. (2019). Cohesin cleavage by separase is enhanced by a substrate motif distinct from the cleavage site. Nature Communications, 10(1), Article 1. 10.1038/s41467-019-13209-y

80. Sato, M., Grant, B. D., Harada, A., & Sato, K. (2008). Rab11 is required for synchronous secretion of chondroitin proteoglycans after fertilization in Caenorhabditis elegans. Journal of Cell Science, 121(Pt 19), 3177–3186. 10.1242/jcs.034678

81. Severson, A. F., & Meyer, B. J. (2014). Divergent kleisin subunits of cohesin specify mechanisms to tether and release meiotic chromosomes. eLife, 3, e03467. 10.7554/eLife.03467

82. Shakes, D. C., Sadler, P. L., Schumacher, J. M., Abdolrasulnia, M., & Golden, A. (2003). Developmental defects observed in hypomorphic anaphase-promoting complex mutants are linked to cell cycle abnormalities. Development (Cambridge, England), 130(8), 1605–1620.

83. Siomos, M. F., Badrinath, A., Pasierbek, P., Livingstone, D., White, J., Glotzer, M., & Nasmyth, K. (2001). Separase is required for chromosome segregation during meiosis I in Caenorhabditis elegans. Current Biology: CB, 11(23), 1825–1835. 10.1016/s0960-9822(01)00588-7

84. Sönnichsen, B., Koski, L. B., Walsh, A., Marschall, P., Neumann, B., Brehm, M., Alleaume, A.-M., Artelt, J., Bettencourt, P., Cassin, E., Hewitson, M., Holz, C., Khan, M., Lazik, S., Martin, C., Nitzsche, B., Ruer, M., Stamford, J., Winzi, M.,…Echeverri, C. J. (2005). Full-genome RNAi profiling of early embryogenesis in Caenorhabditis elegans. Nature, 434(7032), 462–469. 10.1038/nature03353

85. Stein, K. K., & Golden, A. (2018). The C. elegans eggshell. Wormbook, 2018, 1–36. 10.1895/wormbook.1.179.1

86. Stemmann, O., Zou, H., Gerber, S. A., Gygi, S. P., & Kirschner, M. W. (2001). Dual inhibition of sister chromatid separation at metaphase. Cell, 107(6), 715–726. 10.1016/s0092-8674(01)00603-1

87. Sun, Y., Kucej, M., Fan, H.-Y., Yu, H., Sun, Q.-Y., & Zou, H. (2009). Separase is recruited to mitotic chromosomes to dissolve sister chromatid cohesion in a DNA-dependent manner. Cell, 137(1), Article 1. 10.1016/j.cell.2009.01.040

88. Thomas, C., Wetherall, B., Levasseur, M. D., Harris, R. J., Kerridge, S. T., Higgins, J. M. G., Davies, O. R., & Madgwick, S. (2021). A prometaphase mechanism of securin destruction is essential for meiotic progression in mouse oocytes. Nature Communications, 12(1), Article 1. 10.1038/s41467-021-24554-2

89. Tsou, M.-F. B., Wang, W.-J., George, K. A., Uryu, K., Stearns, T., & Jallepalli, P. V. (2009). Polo kinase and separase regulate the mitotic licensing of centriole duplication in human cells. Developmental Cell, 17(3), 344–354. 10.1016/j.devcel.2009.07.015

90. Tunquist, B. J., & Maller, J. L. (2003). Under arrest: Cytostatic factor (CSF)-mediated metaphase arrest in vertebrate eggs. Genes & Development, 17(6), 683–710. 10.1101/gad.1071303

91. Uhlmann, F. (2003). Chromosome Cohesion and Separation: From Men and Molecules. Current Biology, 13(3), R104–R114. 10.1016/S0960-9822(03)00039-3

92. Uhlmann, F., Wernic, D., Poupart, M.-A., Koonin, E. V., & Nasmyth, K. (2000). Cleavage of Cohesin by the CD Clan Protease Separin Triggers Anaphase in Yeast. Cell, 103(3), Article 3. 10.1016/S0092-8674(00)00130-6

93. Viadiu, H., Stemmann, O., Kirschner, M. W., & Walz, T. (2005). Domain structure of separase and its binding to securin as determined by EM. Nature Structural & Molecular Biology, 12(6), Article 6. 10.1038/nsmb935

94. Wainman, A., Giansanti, M. G., Goldberg, M. L., & Gatti, M. (2012). The Drosophila RZZ complex—Roles in membrane trafficking and cytokinesis. Journal of Cell Science, 125(Pt 17), 4014–4025. 10.1242/jcs.099820

95. Waizenegger, I., Giménez-Abián, J. F., Wernic, D., & Peters, J.-M. (2002). Regulation of human separase by securin binding and autocleavage. Current Biology: CB, 12(16), 1368–1378. 10.1016/s0960-9822(02)01073-4

96. Wang, R., Kaul, Z., Ambardekar, C., Yamamoto, T. G., Kavdia, K., Kodali, K., High, A. A., & Kitagawa, R. (2013). HECT-E3 ligase ETC-1 regulates securin and cyclin B1 cytoplasmic abundance to promote timely anaphase during meiosis in C. elegans. Development, 140(10), 2149–2159. 10.1242/dev.090688

97. Watson, E. R., Brown, N. G., Peters, J.-M., Stark, H., & Schulman, B. A. (2019). Posing the APC/C E3 Ubiquitin Ligase to Orchestrate Cell Division. Trends in Cell Biology, 29(2), 117–134. 10.1016/j.tcb.2018.09.007

98. Weber, J., Kabakci, Z., Chaurasia, S., Brunner, E., & Lehner, C. F. (2020). Chromosome separation during Drosophila male meiosis I requires separase-mediated cleavage of the homolog conjunction protein UNO. PLOS Genetics, 16(10), e1008928. 10.1371/journal.pgen.1008928

99. Wessel, G. M., Brooks, J. M., Green, E., Haley, S., Voronina, E., Wong, J., Zaydfudim, V., & Conner, S. (2001). The biology of cortical granules. International Review of Cytology, 209, 117–206. 10.1016/s0074-7696(01)09012-x

100. Wu, J., Larreategui-Aparicio, A., Lambers, M. L. A., Bodor, D. L., Klaasen, S. J., Tollenaar, E., de Ruijter-Villani, M., & Kops, G. J. P. L. (2023). Microtubule nucleation from the fibrous corona by LIC1-pericentrin promotes chromosome congression. Current Biology: CB, 33(5), Article 5. 10.1016/j.cub.2023.01.010

101. Yang, H., Mains, P. E., & McNally, F. J. (2005). Kinesin-1 mediates translocation of the meiotic spindle to the oocyte cortex through KCA-1, a novel cargo adapter. Journal of Cell Biology, 169(3), Article 3. 10.1083/jcb.200411132

102. Yang, H., McNally, K., & McNally, F. J. (2003). MEI-1/katanin is required for translocation of the meiosis I spindle to the oocyte cortex in C. elegans!ZI. Developmental Biology, 260(1), 245–259. 10.1016/S0012-1606(03)00216-1

103. Yu, R., Ren, S.-G., Horwitz, G. A., Wang, Z., & Melmed, S. (2000). Pituitary Tumor Transforming Gene (PTTG) Regulates Placental JEG-3 Cell Division and Survival: Evidence from Live Cell Imaging. Molecular Endocrinology, 14(8), 1137–1146. 10.1210/mend.14.8.0501

104. Zur, A., & Brandeis, M. (2001). Securin degradation is mediated by fzy and fzr, and is required for complete chromatid separation but not for cytokinesis. The EMBO Journal, 20(4), 792–801. 10.1093/emboj/20.4.792

